# Preformed Fibril Seeding Reshapes the Phosphorylated α-Synuclein Proximal Proteome in the Olfactory Bulb

**DOI:** 10.64898/2026.07.22.740026

**Authors:** Solji G. Choi, Atousa Bahrami, Jayda B Duvernay, Tyler Tittle, Ronald Melki, Jeffrey H. Kordower, Bryan A. Killinger

## Abstract

**Background:** Aggregated alpha-synuclein (αsyn) phosphorylated at serine 129 (PS129) accumulates in synucleinopathies, with the olfactory bulb (OB) being severely affected. Non-aggregated physiological PS129 is abundant in the mammalian OB, where it likely modulates αsyn-protein interactions. The impact of aggregation on the PS129 interactome in the OB remains unclear. We <u>hypothesized</u> that αsyn aggregation alters the PS129 interactome, shifting canonical synaptic partners (e.g., SNARE proteins) toward the aggregate-associated network. <u>To test this hypothesis</u>, we mapped PS129 interactions in the OB of PFF-injected WT and SNCA^A53T/A53T^ (M83) mice using biotinylation by antibody recognition (BAR) and pretreated with calf-intestine alkaline phosphatase (CIAP) to distinguish physiological from aggregate-associated PS129 interactomes. Seeding and spread were assessed by immunohistochemistry and in situ seeding immunodetection assay (isSID).

**Results:** Following OB-PFF injections, CIAP-resistant aggregates and seeds were detected throughout the neuroaxis of M83 mice (e.g., OB to brainstem) but less so in WT mice. isSID seeding was concentrated near CIAP-resistant aggregates, but the overlap was only partial. BAR-PS129 identified 2,309 proteins in M83 OBs and 990 proteins in WT OBs. Of these, 357 proteins in M83 mice and 247 proteins in WT mice were associated with the CIAP-resistant, aggregate-enriched PS129 fraction. In both models, the CIAP-resistant interactome largely overlapped with the broader PS129 interactome, suggesting that seeded aggregation primarily affects existing PS129 interactions rather than directing the formation of new pathological ones. A conserved 107-protein CIAP-resistant signature shared between M83, and WT mice was enriched for axon–glia adhesion, myelin-associated, axonal/cytoskeletal, proteostatic, and synaptic vesicle-related proteins.

**Conclusion:** Aggregated PS129 engages a subset of PS129 networks enriched at axon–glial interfaces. CIAP-resistant aggregates were partially associated with seed competency, indicating that CIAP resistance and seed competency are related but not equivalent. These results provide molecular details of αsyn seeding and spread from the OB.

## Introduction

Synucleinopathies, including Parkinson’s disease (PD) and dementia with Lewy bodies (DLB), are neurodegenerative diseases defined by intraneuronal α-synuclein (αsyn) aggregates, collectively termed Lewy pathology (LP). αSyn aggregation may begin in the olfactory bulb (OB), which then spreads through connected circuits via a seeding-and-spread mechanism [1–3]. Systematic histopathological studies have noted that the OB consistently bears LP, even in individuals without a clinical diagnosis [1–3]. Although LP is detected throughout the OB, the deepest layers, including the granule cell and inner plexiform layer, are most severely affected [1]. OB-LP occurs predominantly in neurons but can also be detected in glial cells, including astrocytes [4]. Despite the importance of OB as a potential start site for synucleinopathies, the precise mechanism underlying αsyn aggregation in the OB remains unclear.

Injection of αsyn fibrils made *in vitro* (preformed fibrils, “PFFs”) into the wild-type (WT) mouse OB (“OB-PFF”) has been used to model seeding/spreading mechanisms [5–11]. OB-PFF mice develop proteinase K-resistant aggregates at the injection site (i.e., the granule cell layer of the OB), as well as the anterior olfactory nucleus, and interconnected cortical brain regions such as the piriform cortex [5, 8–10]. However, it’s still unclear whether the progressive spread occurs to the midbrain or the brainstem [7]. αSyn spreads from the OB via retrograde transport along centrifugal OB projections, but anterograde transport along mitral cell axons is also likely [4, 6]. OB-PFF injections induce acute neuronal loss in the deep layers of the OB but spare mitral cells [5].

Phosphorylated αsyn at serine 129 (PS129) is a widely used marker of αsyn aggregates, but it also occurs under physiological conditions in response to neuronal activity, where it promotes protein interactions at synaptic terminals [12–15]. We recently discovered that aggregated-PS129 was phosphatase resistant, abundant in well-preserved brain specimens (i.e., lacking postmortem interval), and that calf intestinal alkaline phosphatase (CIAP) could be used to distinguish pathological from physiological PS129 [15, 16]. This discovery was especially important in the OB and in animal models, where physiological PS129 is abundant [7, 13, 14]. Although physiological PS129 is largely undetectable in postmortem human brain because of rapid postmortem dephosphorylation [15, 16], physiological PS129 is readily detected in rapidly fixed tissues (e.g., perfusion fixed rodent brain and human surgical specimens) [13, 14]. The role of PS129 in disease remains unclear, with evidence supporting protective [17, 18], toxic [19, 20], or inconsequential effects [14, 21, 22]. Nevertheless, normal αsyn-binding partners, such as VAMP2, are sequestered within LP [14, 23–25] raising two questions: Does aggregation sequester αsyn’s normal partners and impair their function, resulting in a loss of function, or does it promote aberrant interactions that drive a toxic gain of function?

The impact of aggregation on the PS129 protein environment (proximal proteome) has been difficult to resolve because abundant physiological PS129 can obscure aggregate-specific interactions. Our group has pioneered the use of biotinylation by antibody (BAR) recognition to measure in situ αsyn interactomes or “proximal proteomes” (i.e., spatial interactomes) [13, 15, 26–28], particularly in the human brain. However, previous efforts in OB-PFF mice were similarly limited by physiological PS129 [7]. The influence of aggregation on PS129 interactions is critical but remains uncharacterized in the brain. Although BAR has been used to characterize physiological and pathological PS129 proximal proteomes, as well as the total αsyn interactome, in human and mouse brain [7, 13, 15, 27, 28], it has not been applied to directly determine how aggregation alters the PS129 proximal proteome within the same brain region in vivo.

We hypothesize that αsyn aggregation induces a pathological shift in the PS129 interactome in the OB, moving away from its normal synaptic interaction partners toward a disease-associated network that may promote the spread of αsyn pathology to anatomically connected regions. To test this, we injected PFFs into the OBs of WT mice [5, 7, 10, 29] and A53T αsyn transgenic mice (M83). Physiological and aggregated PS129 were distinguished within the same OB using CIAP pretreatment [15, 16] and proximal proteins were mapped using BAR.

## Materials and Methods

### Animals and Tissue Processing

Male and female C57BL/6J mice (WT; RRID: IMSR_JAX:000664; n = 5 per treatment condition) and homozygous B6;C3-Tg(Prnp-SNCA*A53T)83Vle/J mice (M83; RRID: IMSR_JAX:004479; n = 21 per treatment condition) were used for all studies [5, 7–10, 30, 31]. When the oldest M83 mice reached 8 months of age, all animals were euthanized with CO_2_ and immediately perfused with PBS followed by 4% paraformaldehyde (PFA), as previously described [16]. Whole brains were then removed and post-fixed in 4% PFA at 4°C for over 24 hours. Brains were subsequently equilibrated in 15% and then 30% sucrose, frozen, and sectioned at 40 μm on a freezing-stage microtome. Sections were stored at −20°C in cryoprotectant solution (30% sucrose and 30% ethylene glycol in PBS) [27]. All animal procedures were reviewed and approved by the Rush University Medical Center Institutional Animal Care and Use Committee.

### αsyn Preformed Fibrils

Human αsyn PFFs were amplified in the presence of dilute frontal cortex homogenates from individual with PD as previously described [7, 32]. Prior characterization of fibril assembly, morphology, and biophysical properties showed that the amplified PFFs were similar and induced comparable pathological spread and behavioral deficits after OB injection in WT mice [7]. Amplified PFF aliquots were sonicated, frozen, and stored at −80°C until use [7, 33].

### Stereotactic Injection

M83 and WT mice received unilateral stereotactic injections of human αsyn PFFs or PBS into the OB granule cell layer. Human αsyn PFFs were injected at 5 μg/μL, with a total injection volume of 2 μL using the following coordinates relative to bregma and the dural surface: AP, +5.4 mm; ML, -0.75 mm; DV, −1.0 mm. Injections were performed at 0.2 μl/min using a 10 μl Hamilton microsyringe fitted with a 30-gauge needle [7, 9, 10]. After injection, the needle was left in place for 5 min and then slowly withdrawn [7, 9]. Before injection, PFF aliquots were thawed at 37°C for 3 min, briefly spun down, and immediately transferred to the syringe. Mice were maintained for approximately 100 days after PFF or PBS injection before euthanasia.

### Calf Intestinal Alkaline Phosphatase (CIAP) Treatment

The CIAP treatment protocol was performed as described in protocols.io (https://dx.doi.org/10.17504/protocols.io.5qpvoky7bl4o/v3) and in our previous study [16]. Free-floating sections were washed three times in dilution medium (DM; 50 mM Tris-HCl, pH 7.2, 150 mM NaCl, 0.05% Triton X-100) and then incubated in 1% Triton X-100 in DM for 10 min. Sections were rinsed three times in DM and subsequently washed twice in CIAP buffer (100 mM NaCl, 10 mM MgCl₂ , and 50 mM Tris-HCl, pH 7.9). Sections were then incubated in CIAP buffer containing 0.06 U/µL calf intestinal alkaline phosphatase (Promega, Catalog#: M1821) for 24-28 h at 37°C. After incubation, sections were washed in DM and subjected to heat-induced antigen retrieval (HIAR) in sodium citrate buffer (10 mM sodium citrate, 0.05% Tween 20, pH 6.0) at 85°C for 30 min, followed by cooling to room temperature.

### Immunohistochemistry

After pretreatment with or without CIAP, free-floating mouse brain sections were rinsed in DM and incubated for 1 h at room temperature in blocking/quenching buffer containing 3% goat serum, 2% BSA, 0.4% Triton X-100, 0.3% hydrogen peroxide, and 0.1% sodium azide in DM. Sections were then washed in DM and incubated overnight with anti-PS129 antibody (Abcam, EP1536Y; 1:50,000 in blocking buffer; RRID:AB_869973). The following day, sections were washed in DM and incubated for 1 h at room temperature with HRP-conjugated goat anti-rabbit secondary antibody (Thermo Fisher Scientific, Cat# 31462; 1:1,000 dilution; RRID:AB_228338) in DM containing 1% BSA and 1% goat serum. Sections were then rinsed in DM followed by sodium acetate buffer (0.2 M imidazole, 1.0 M sodium acetate, pH 7.2).

Immunoreactivity was visualized using a standard nickel-enhanced 3,3′-diaminobenzidine (DAB)-imidazole reaction. Sections were subsequently washed in sodium acetate buffer and PBS, then mounted onto glass slides. Finally, sections were counterstained with methyl green, dehydrated, cleared in xylenes, and coverslipped with Cytosol XYL (Epredia, 8312-4).

### In Situ Seeding Immunodetection Assay

Procedures were adapted from [34]. Free-floating sections were washed in TBS (50 mM NaCl, 50 mM Tris-HCl, pH 7.6) and quenched in 0.3% hydrogen peroxide in TBS for 30 min at room temperature. Sections were then subjected to HIAR in citrate buffer (10 mM citric acid, 0.05% Tween 20, pH 6.0) for 10 min at 95–100°C. Sections were rinsed in PBS and then in PIPES buffer (100 mM, pH 6.9), followed by preconditioning in PIPES buffer for 1 h. Sections were incubated overnight at 37°C with recombinant His-tagged mouse αsyn monomer (ACRO Biosystems; Cat# ALN-M52H6) diluted in PIPES buffer to a final concentration of 0.025 mg/ml. After incubation, sections were washed in TBS and fixed in 4% PFA for 10 min, then washed in PBS containing 0.5% Tween 20 (PBS-T). Sections were blocked in 10% normal horse serum with 0.5% Triton X-100 containing TBS for 1 h at room temperature. Sections were then incubated for 2 h at room temperature with an HRP-conjugated anti-6×His antibody (Thermo Fisher Scientific; Cat# MA1-21315-HRP) diluted 1:5,000 in blocking buffer.

Sections were washed twice in PBS-T for 5 min, and His-tagged αsyn immunoreactivity was visualized using either nickel-enhanced DAB or tyramide fluorescence labeling. A detailed protocol is available at https://dx.doi.org/10.17504/protocols.io.261gey7bov47/v2.

Multiplex Tyramide Signal Amplification Labeling Dual labeling of isSID and CIAP-resistant PS129 was performed as follows. After the isSID, tissue sections were rinsed in borate buffer (0.05 M sodium borate, pH 8.5), and signal was developed for 30 min in tyramide fluorophore solution containing 0.003% hydrogen peroxide and 5 μM CF568 (Biotium) in borate buffer (Fisher Scientific; Cat# 50-196-4912). After CF568 labeling, sections were treated with CIAP and heat-induced antigen retrieval (HIAR) in sodium citrate buffer. Sections were then quenched and blocked for 1 h in goat serum-containing blocking buffer, incubated overnight at 4°C with PS129 primary antibody, washed in DM, and incubated for 1 h with HRP-conjugated anti-rabbit secondary antibody. Sections were rinsed in DM followed by borate buffer and developed for 30 min in CF488 tyramide fluorophore solution (5 μM; Biotium, Cat# NC1474216).

GFAP and CIAP-resistant PS129 labeling was performed as follows. Sections first underwent CIAP treatment and HIAR in sodium citrate buffer, followed by quenching and blocking for 1 h at room temperature in DM containing 0.3% hydrogen peroxide, 0.1% sodium azide, 3% horse serum, 2% bovine serum albumin, and 0.4% Triton X-100. Sections were incubated overnight at 4°C with anti-GFAP primary antibody (Abcam, EPR1034Y; RRID: AB_3669042), washed in DM, and incubated for 1 h at room temperature with peroxidase-conjugated horse secondary antibody (Vector Laboratories; SKU# PI-9500-1; 1:1,000) diluted in DM containing 1% horse serum and 1% BSA. Sections were washed in DM and borate buffer and developed for 30 min at room temperature in borate buffer containing 5 μM CF568. Sections were then washed, subjected to HIAR, cooled, and labeled for PS129 with CF488 tyramide fluorophore as described above.

After dual labeling, all sections were rinsed in PBS, stained with DAPI (Invitrogen; Cat# D1306; 1:2,000) for 30 min, washed again in PBS, mounted onto slides, coverslipped with antifade mounting medium (Abcam; ab104135), and sealed with nail polish.

### Microscopy and Imaging

For brightfield imaging, prepared slides were scanned using a Li-Cor Odyssey M system at 5 μm resolution with Li-Cor acquisition software. Whole-slide brightfield scans were imported into Adobe Photoshop for downsampling, cropping, and automated color and brightness adjustment. Higher-magnification brightfield images were acquired on a Nikon A1R inverted confocal microscope equipped with a transmitted-light brightfield detector, using 20× or 60× objectives.

For multiplex fluorescence imaging, confocal images were acquired using a Nikon A1R inverted confocal microscope. Whole-tissue scans were acquired as large z-stacks using a 10× objective, denoised, and processed as maximum-intensity projections for presentation. Higher-magnification confocal images were acquired using 20× or 60× objectives and deconvolved in NIS-Elements software. All images were assembled in Adobe Illustrator for figure layout and final presentation.

### Quantification of PS129 Immunoreactivity

Quantification of the PS129 signal across brain regions was performed in NIS-Elements using RGB-based thresholding and the Macro function, as previously described [16, 28, 35]. For each brain region, 20× images were acquired within an 800 × 800-pixel bounding box. RGB color thresholds were manually adjusted to exclude methyl green nuclear staining, which did not overlap with the black/purple PS129 signal, and the same threshold was batch-applied to all images regardless of treatment group or CIAP condition. The resulting binary area fraction was exported to Excel and expressed as a percentage by multiplying the raw values by 100. Representative quantified images are shown in Fig. S1. Statistical comparisons between treatment conditions within each brain region were performed using two-tailed Mann–Whitney U tests. P values were corrected for multiple comparisons across brain regions using a false discovery rate of 5% with the two-stage step-up method of Benjamini, Krieger, and Yekutieli [36].

### Biotinylation by Antibody Recognition (BAR)

BAR-PS129 was performed on 15–20 pooled M83 OBs per capture to generate the four interactomes shown in Figure 1A, with equal numbers of bulbs used across conditions within each case (Supplementary Table 1; mean wet mass: 3.617 ± 1.757 mg). The same BAR-PS129 protocol was used for WT mice (Fig. 5A), with an average pooled OB wet mass of 4.51 ± 0.79 mg across all BAR-capture samples (Supplementary Table 1).

**Figure 1.**
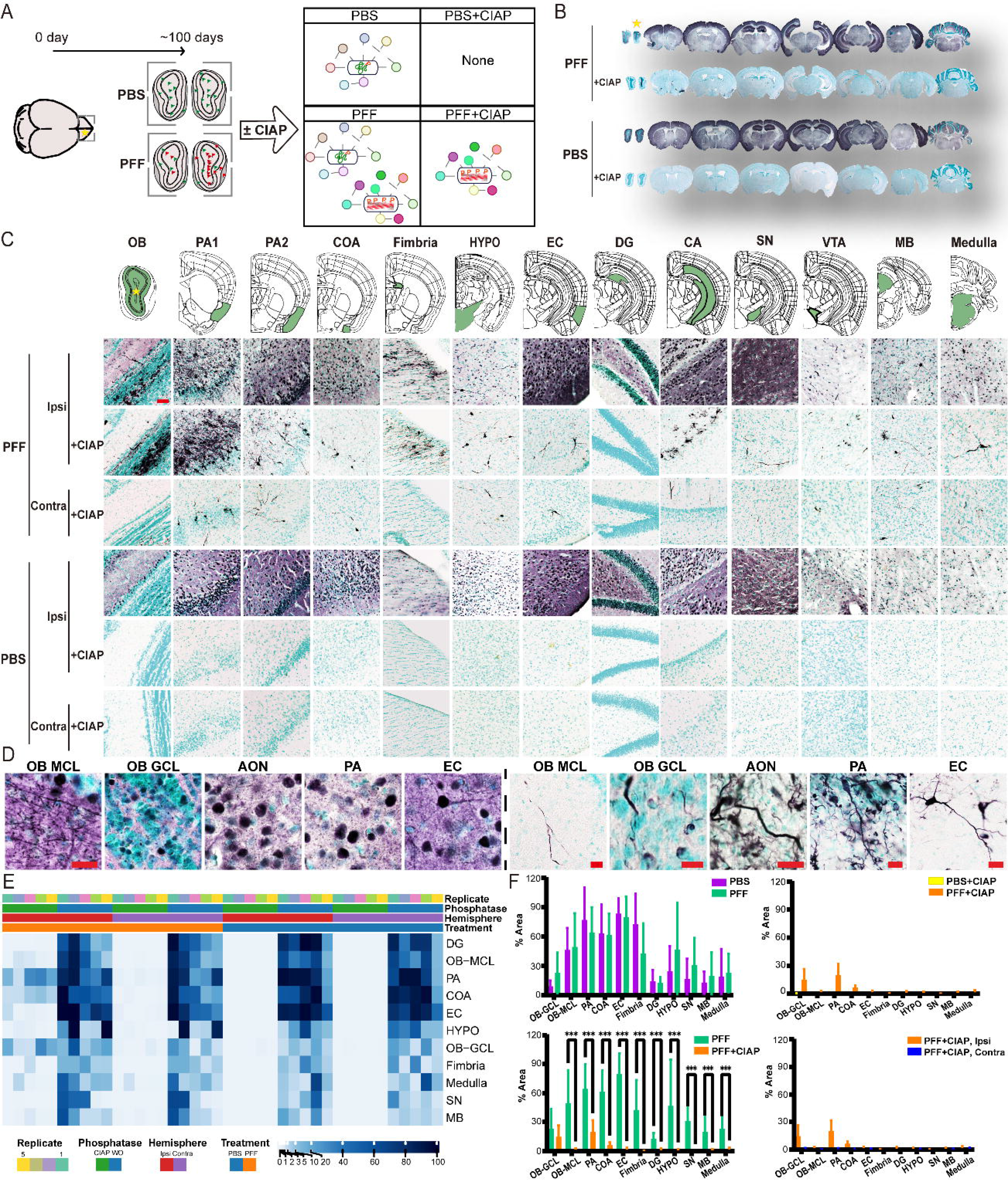
Seeding and spread following PFF injection in the OB of M83. (A) Experimental design for detecting distinct PS129 species and their protein–protein interactomes in the M83-PFF model. M83 mice received OB injections of PFFs or PBS and were transcardially perfusion-fixed 100 days later. Coronal brain sections were pretreated with or without CIAP, followed by BAR to measure PS129-associated interactomes. Comparison of CIAP-treated and untreated conditions enabled differentiation of aggregate-associated and non-aggregate-associated PS129 interactions. (B) PS129 IHC staining across the neuroaxis of PBS- and PFF-treated M83 mice, with or without CIAP pretreatment. The star indicates the PFF injection site. (C) Distribution of CIAP-resistant PS129 across the neuroaxis of PBS- and PFF-treated M83 mice. Regions containing CIAP-resistant PS129 are annotated in green on reference atlas sections, with representative images shown for each annotated region. Atlas images were adapted from the Allen Institute Mouse Brain Reference Atlas. Scale bar, 50 μm. (D) High-magnification images of CIAP-sensitive and CIAP-resistant PS129 in five brain regions of M83 mice. Scale bars, 20 μm for CIAP-sensitive PS129 in all five regions; 10 μm for CIAP-resistant PS129 in the OB GCL; and 20 μm for CIAP-resistant PS129 in all other regions. (E, F) Heatmap and bar-graph quantification of PS129-positive area across 11 brain regions containing CIAP-resistant PS129. Statistical comparisons were performed using two-tailed Mann–Whitney U tests with FDR correction. ***q < 0.001. n=5 animals per region. Abbreviations: OB, olfactory bulb; MCL, mitral cell layer; GCL, granular cell layer; PA1, piriform area proximal to the OB; PA2, piriform area distal to the OB; COA, cortical amygdalar area/nucleus of the lateral olfactory tract; HYPO, hypothalamus; EC, entorhinal cortex; DG, dentate gyrus; CA, cornu ammonis; SN, substantia nigra; VTA, ventral tegmental area; MB, midbrain, including the periaqueductal gray, superior colliculus, and motor-related deep gray/white layers.

Free-floating sections were labeled in situ with anti-PS129 primary antibody (Abcam, EP1536Y) followed by HRP-conjugated secondary antibody. Sections were incubated for 30 min at room temperature in borate buffer containing biotinylated tyramide (14.7 μL of 12.5 mg/mL stock in DMSO; Sigma-Aldrich, SML2135) and 0.003% hydrogen peroxide (Sigma-Aldrich, H1009), washed in PBS, and solubilized in crosslink-reversal buffer containing 5% SDS, 500 mM Tris-HCl pH 7.6, 150 mM NaCl, 2 mM EDTA pH 8.0, and 1 mM PMSF. Samples were briefly sonicated, vortexed, heated at 98°C for 30 min, and centrifuged at 22,000 × g for 30 min.

Supernatants were diluted in TBST and incubated with 40 μL streptavidin-coated magnetic beads (Thermo Fisher Scientific; Cat# PI88817) for 1 h at room temperature with gentle rotation. Beads were magnetically isolated, washed twice for 30 min in wash buffer containing 150 mM NaCl, 50 mM Tris-HCl pH 7.6, 1 mM EDTA pH 8.0, 0.1% SDS, and 1% Triton X-100, followed by a 16 h wash at 4°C. Captured proteins were eluted at 98°C for 10 min in 40 μL LDS sample buffer (Invitrogen; B0008) containing 5 mM DTT (Roche; 10708984001).

For LC-MS/MS preparation, 30 μL of each eluate was briefly electrophoresed into 4–12% Bis-Tris gels (Fisher Scientific; NW04127BOX) at 200 V, fixed in 50% ethanol/10% acetic acid, rehydrated, stained with Coomassie blue (Invitrogen; LC6060), and rinsed to remove excess stain. The entire lane was excised and submitted for trypsin digestion and LC-MS/MS analysis.

### Liquid Chromatography–Tandem Mass Spectrometry (LC–MS/MS) Analysis

Tryptic peptides were analyzed on an Orbitrap Exploris 240 mass spectrometer. M83 samples were analyzed in duplicate LC-MS/MS injections, while WT with single injections. Raw spectra were processed in MaxQuant and searched against the reviewed Swiss-Prot canonical mouse proteome (UP000000589). Label-free quantification was performed using MaxLFQ, which normalizes MS1 intensities across runs using a sample-similarity network to improve relative quantification and reduce missing values [37, 38]. Duplicate technical-injection values for each M83 sample were averaged in R before downstream analysis.

### The resulting proteinGroups.txt output was analyzed in LFQ-Analyst [39]

Differential abundance was assessed using protein-wise linear models with empirical Bayes moderation and Benjamini–Hochberg correction, applying an adjusted p-value threshold of <0.05 and an absolute log₂ fold-change cutoff of 1 [40]. Single-peptide identifications were excluded [41, 42]. Proteins detected in each BAR condition were also ranked using an importance score, defined as log₂ fold change × −log₁₀ adjusted p value, to integrate effect size with statistical confidence [43, 44]. Full LFQ-Analyst outputs and calculated importance scores for all proteins identified in both animal models are provided in Datasets S1–S3.

### Spot Blotting Analysis

1 μL of BAR bead eluate was applied to a methanol-activated PVDF membrane and air-dried. Membranes were reactivated in methanol, rinsed in ultrapure water, fixed in 4% PFA for 30 min, washed in TBST, and blocked for 1 h at room temperature in 5% BSA/TBST. Biotinylated proteins were detected by incubating membranes with ABC reagent (Vector Laboratories) diluted in blocking buffer for 30 min at room temperature.

For αsyn detection, membranes were blocked in 5% nonfat dry milk/TBST and incubated overnight at 4°C with SYN1 antibody (BD Biosciences; #610787; RRID: AB_398108; 1:2,000). Membranes were washed in TBST and incubated for 1 h with HRP-conjugated anti-mouse secondary antibody (1:6,000; RRID: AB_330924).

Membranes incubated for ABC or αsyn were developed using Clarity Western ECL substrate (Bio-Rad; #170-5060) and imaged on a ChemiDoc system (Bio-Rad; RRID: SCR_019037). For quantification, images were acquired using optimal auto-exposure, exported as raw TIFF files, and analyzed in ImageJ (v1.54g; RRID: SCR_003070). Data were plotted and analyzed in GraphPad Prism 10 using two-way ANOVA with uncorrected Fisher’s LSD.

### Visualization of BAR-Enriched Proteins

Plots showing protein identification counts, heatmaps, and PCA were generated in LFQ-Analyst from MaxQuant LFQ output and imported for figure preparation. Proteins with calculated importance scores were imported into GraphPad Prism 10, plotted, and compared across treatment groups using the Kruskal–Wallis test followed by Dunn’s multiple comparisons test. All other graphs were generated in R (v4.4.1) using EnhancedVolcano, readxl, dplyr, ggplot2, ggrepel, patchwork, stringr, RColorBrewer, and wordcloud for data handling and visualization, including volcano plots, top 20 importance score-ranked proteins, and top 20 Gene Ontology term clouds identified using g:Profiler [45]. Protein–protein interaction networks were visualized and annotated in Cytoscape (v3.10.3) using stringApp (v2.2.0) and CentiScaPe (v2.2) [46–48].

## Results

### CIAP-resistant PS129 reveals αsyn aggregate spread in M83 mice

M83 mice received unilateral OB injections of PBS or PFFs and were euthanized 100 days later. Coronal sections were treated ±CIAP and labeled by BAR-PS129 (Fig. 1A). The model was characterized before BAR-PS129 data analysis. Without CIAP, PS129 immunostaining was widespread in PBS- and PFF-treated M83 brains (Fig. 1B). CIAP abolished PS129 reactivity in PBS treated M83 mice, whereas CIAP-resistant PS129 remained in PFF-treated mice, mostly at the OB-GCL injection site and downstream brain nuclei (cortical amygdalar area (COA) and piriform area (PA), entorhinal cortex (EC), hippocampal formation, midbrain (MB), and medulla. In PFF-treated mice, bilateral CIAP-resistant PS129 appeared as rounded cytoplasmic labeling across multiple regions, accompanied by dense, thread-like neuritic structures resembling Lewy neurites, some extending from an enlarged, abnormal perikaryon (Fig. 1C). In the OB, PS129 staining was restricted to the GCL, inner plexiform layer, and to a limited extent, outer plexiform layer. Mitral cells and their apical dendrites were strongly PS129-reactive in PBS- and PFF-injected mice [13, 14, 49] (Fig. 1D, left panels). The AON, PA, and EC also showed dense PS129 puncta, cell bodies, and neurites. In contrast, in the same brain regions of PFF-treated M83 mice, CIAP-resistant PS129 (Fig. 1D, right panels) appeared as intracellular coil-like tangles (i.e., coiled bodies) and swollen dystrophic neurites.

PS129-positive area was quantified across 11 atlas-defined brain regions. The data are presented as a clustered heatmap (Fig. 1E) and a statistical analysis (Fig. 1F). Without CIAP, PBS- and PFF-injected M83 mice showed no significant regional differences in the ipsilateral PS129-positive area. CIAP markedly reduced the PS129 signal in both groups, with residual signal detected mainly in the GCL, PA, and COA. Direct comparison of ipsilateral PFF and PFF+CIAP sections showed significantly higher PS129-positive area in PFF tissues across 10 of 11 regions, with the GCL injection site as the only exception (Fig. 1F, bottom right; ***q < 0.001, FDR-corrected Mann–Whitney U tests). Together, results show that following CIAP, robust seeding and spread throughout the neuroaxis of OB-PFF M83 mice.

### CIAP-resistant PS129 reveals limited αsyn aggregate spread in WT mice

WT mice showed similar PS129 staining in the PFF and PBS groups (Fig. 2A), with immunoreactivity distributed throughout the neuroaxis in both, consistent with previous reports [7, 13, 16]. CIAP treatment abolished PS129 signals, leaving only residual CIAP-resistant labeling at the injection site, marked by a star in Fig. 2A & C. In the contralateral OB, PS129 labeling was sparse, with only rare CIAP-resistant rod-like structures (Fig. 2B). In the AON, which projects predominantly to the deep OB GCL bilaterally [50, 51], aggregated PS129-positive somatic and neuritic inclusions were detected in both hemispheres (Fig. 2B). After CIAP treatment, dysmorphic neurites and occasional somatic inclusions persisted bilaterally, with greater pathology ipsilaterally (Fig. S2A). In PA, PS129 was diffuse, whereas CIAP-resistant pathology was limited to occasional neurites and absent contralaterally (Fig. 2B & S2B). In PFF-treated WT mice, dysmorphic neurites appeared at the OB injection site and persisted after CIAP (Fig. 2C). Post-CIAP, PS129 was mainly absent outside the OB injection area. PBS-treated WT mice showed a complete loss of PS129 after CIAP.

**Figure 2.**
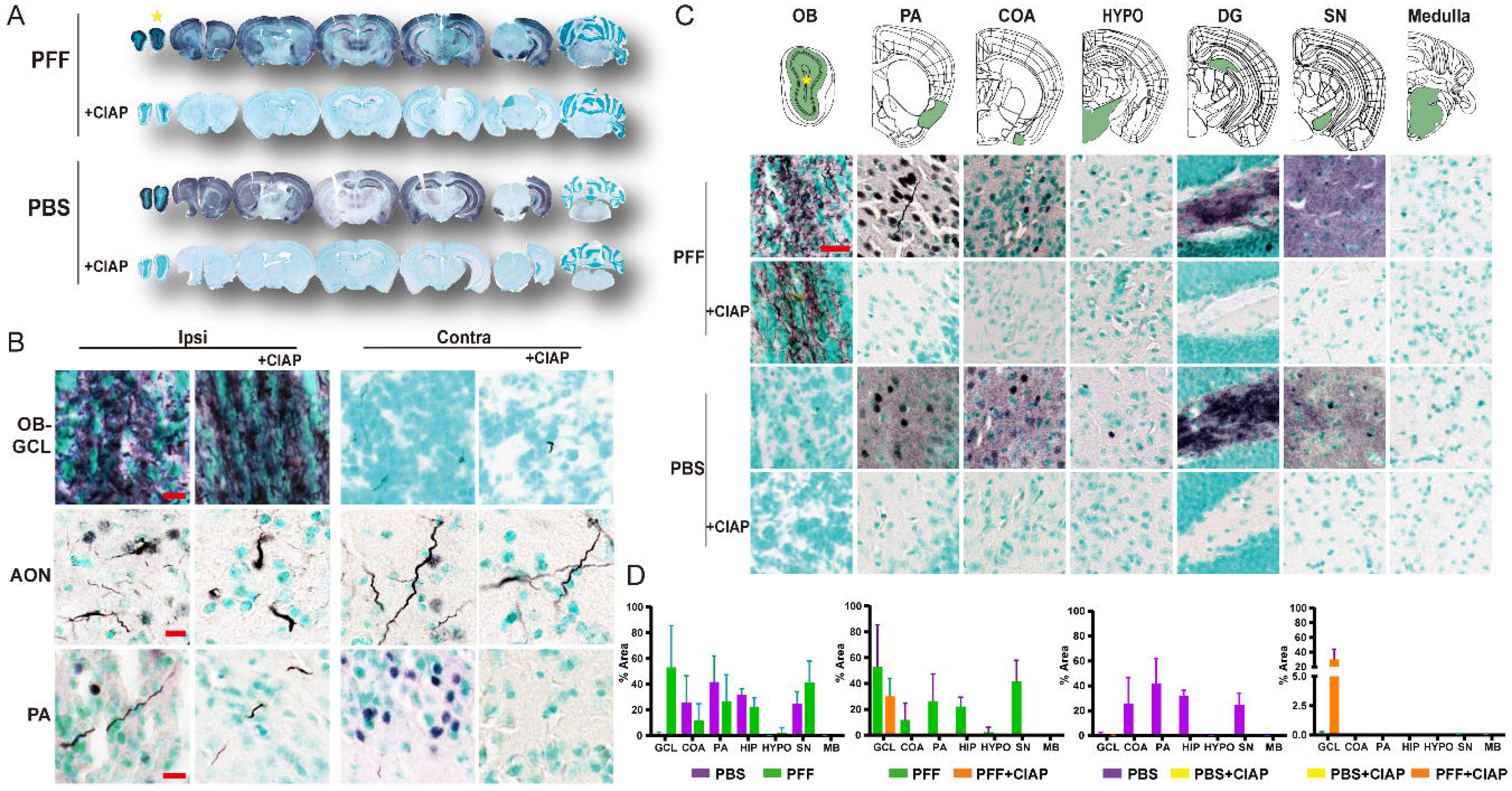
Seeding and spread following PFF injection in the OB of WT. WT mice were processed in parallel with M83 mice, from OB injection through histological characterization. (A) PS129 IHC staining across the neuroaxis of PBS- and PFF-treated WT mice, with or without CIAP pretreatment. The star indicates the PFF injection site. (B) Enlarged 20× images or 60x images of the OB GCL, anterior olfactory nucleus (AON), and PA from both hemispheres, with and without CIAP treatment. Scale bars, 10 μm for all images. (C) Ipsilateral distribution of PS129 staining across brain regions highlighted in green on the atlas for each condition. Scale bar, 25 μm. (D) Bar graphs showing quantification of PS129-positive area across seven brain regions under each condition. Statistical comparisons were performed using FDR-corrected Mann–Whitney U tests; no significant differences were detected. n=3.

Quantification across the seven atlas-defined regions confirmed that PBS- and PFF-treated WT mice had similar overall PS129 immunoreactivity (Fig. 2D, no significant differences were detected by FDR-corrected Mann–Whitney U tests). After CIAP treatment, a detectable PS129 signal remained primarily in the GCL of PFF-treated mice, whereas all other regions showed minimal or undetectable immunoreactivity. Together, these findings indicate that CIAP-resistant PS129 pathology in WT mice was mostly confined to the injection site.

### CIAP-resistant PS129 pathology coincides with seeding propensity

αsyn seeding is required for sustained, progressive spreading [31, 52]. To test whether CIAP-resistance corresponded with seeding, we performed an isSID in both WT and M83 mice. Seed-competent species were detected exclusively in brain regions containing CIAP-resistant PS129 aggregates of OB-PFF-injected M83 mice (Fig. S3A, B), whereas WT brains had a weak isSID signal restricted to the OB-PFF injection site (i.e., PFF or PBS) (Fig. S3C). Multiplex tyramide-fluorophore labeling of isSID and CIAP-resistant PS129 in M83-PFF tissue showed dense accumulation of CIAP-PS129 and seeds (Fig. 3A). However, isSID appeared as irregular granular or globular deposits, whereas CIAP-resistant PS129 localized mainly to neuritic structures and intracellular inclusions. Both isSID and CIAP-PS129 were lower in the contralateral hemisphere but were detected bilaterally. The partial overlap of isSID and CIAP-PS129 was also present in the piriform area and NLOT/COA, with stronger ipsilateral labeling. isSID labeled the midbrain, especially the periaqueductal gray, and the medulla, where both signals appeared in neuritic and somatic aggregates.

**Figure 3.**
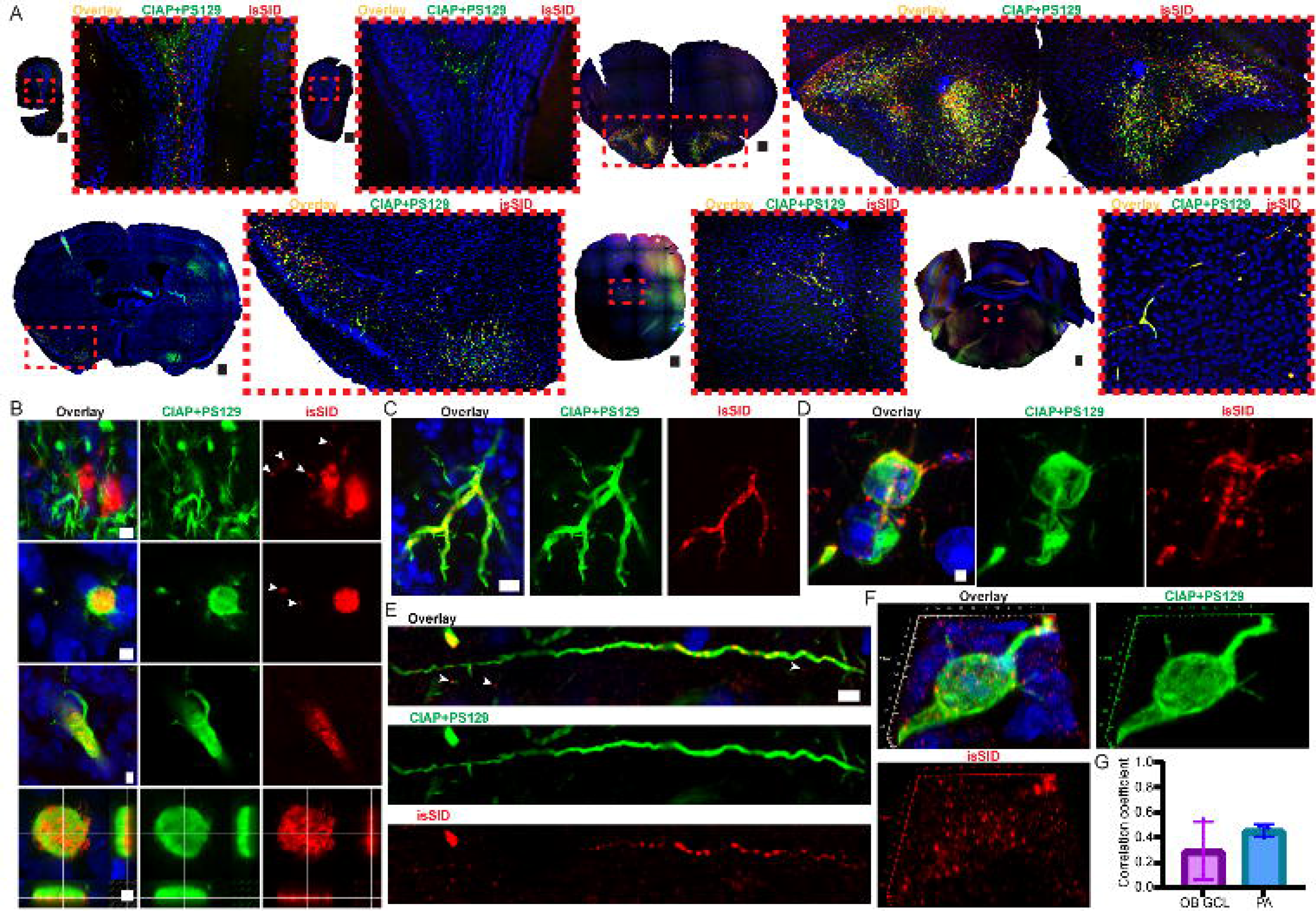
Spread of CIAP-resistant PS129 and seed-competent αsyn species following PFF injection into the OB of M83 mice. M83 mice injected in the OB with PFFs were analyzed by multiplex tyramide signal amplification labeling for CIAP-resistant PS129 and isSID. (A) Whole-tissue z-stack maximum-intensity projections show dual labeling across multiple brain regions. Red boxes indicate enlarged regions. Top panels show ipsilateral OB, contralateral OB, and AON-containing regions; bottom panels show PA/NLOT/COA-containing regions, midbrain containing the periaqueductal gray, and medulla-containing regions (Hindbrain; HB). The left hemisphere is ipsilateral in all panels. Scale bars, 300 μm. (B) Higher-magnification images of the ipsilateral OB GCL. The first row shows globular isSID structures that are morphologically distinct from CIAP-resistant PS129-positive neuritic pathology. The second and third rows show Lewy body– or Lewy neurite–like PS129-positive structures containing isSID deposits, with surrounding punctate isSID signal indicated by white arrows. The last row shows an orthogonal view of a Lewy body–like aggregate with isSID-positive patches within and around the PS129-positive structure. Scale bars, 5 μm for the top 3 rows, and 2.5ul for the orthogonal view, respectively. (C) OB external plexiform layer-to-glomerular layer region showing isSID-positive puncta along a subset of CIAP-resistant PS129-positive dystrophic neurites. Scale bar, 10 μm. (D) AON intracellular CIAP-resistant PS129-positive structure with punctate isSID signal within the soma and along extended neuritic processes. White arrows indicate nuclear fragments within an isSID-associated neuritic structure; see Fig. S4C for the nuclear channel. Scale bar, 2 μm. (E) Elongated CIAP-resistant PS129-positive neuritic process in the AON containing punctate isSID signal, with additional puncta surrounding the process. White arrows indicate punctate structures. Scale bar, 5 μm. (F) Three-dimensional reconstruction of a CIAP-resistant PS129-positive Lewy body–like aggregate with extended processes in the PA, showing abundant isSID signal within and around the aggregate. (G) Pearson’s correlation coefficients between isSID and CIAP-resistant PS129 signals in the OB GCL and PA. Mean coefficients were 0.2949 in OB GCL and 0.4551 in PA, with SD = 0.2304 and 0.04842, respectively. n = 3 mice.

isSID and CIAP-resistant PS129 were closely associated in the ipsilateral OB GCL (Fig. 3B, Fig. S4A). isSID typically persisted as large globular deposits and smaller punctate structures that were morphologically distinct from the dystrophic neurites labeled by CIAP-resistant PS129. Occasional overlap was also observed in somatic structures and swollen neuritic processes. In neuritic structures, small isSID-positive puncta were detected within CIAP-resistant PS129-positive processes, with additional puncta surrounding these neurites (Fig. S4A). Orthogonal imaging of a Lewy body-like structure further revealed discrete isSID-positive patches within and around dense PS129-positive structures (Fig. 3B, bottom panels).

Additional OB regions, including the EPL-to-GL region, showed occasional CIAP-resistant PS129-positive neuritic pathology (Fig. 3C). In this region, CIAP-resistant PS129 labeled elongated dystrophic neurites, whereas isSID appeared as discrete puncta along PS129-positive neurites. In the AON, where both signals were strongest, CIAP-resistant PS129 labeled large perinuclear inclusions, with isSID closely associated and prominent along connected neurites. (Fig. 3D). Punctate isSID along CIAP-PS129 neurite was also observed in the AON (Fig. 3E), with similar patterns observed in the PA (Fig. S4B). Three-dimensional reconstruction of a CIAP-resistant PS129-positive Lewy body-like inclusion with extended processes further showed abundant punctate isSID signal within and around the PS129-positive structure, with partial overlap between the two signals (Fig. 3F and Fig. S4B). In the OB GCL and PA, the two signals showed moderate colocalization (Fig. 3G). Together, CIAP-PS129 is spatially related to, not equivalent to, αsyn seeds.

### CIAP-resistant PS129 proximal proteome in M83 OBs

BAR-PS129 labeling is shown in Figure 4A (Fig. S5). All non-CIAP conditions showed strong enrichment when compared to BAR-Neg (>14-fold for biotin and >120-fold for αsyn), whereas CIAP markedly reduced enrichment (Fig. 4B, C). αSyn enrichment was lower in PFF-treated than PBS-treated OBs (156.2-fold vs. 208.7-fold; p = 0.0071). After CIAP, αsyn enrichment was higher in PFF-treated than PBS-treated OBs (5.43-fold vs. 0.7804-fold), indicating successful BAR capture of CIAP-resistant PS129 in PFF OBs.

**Figure 4.**
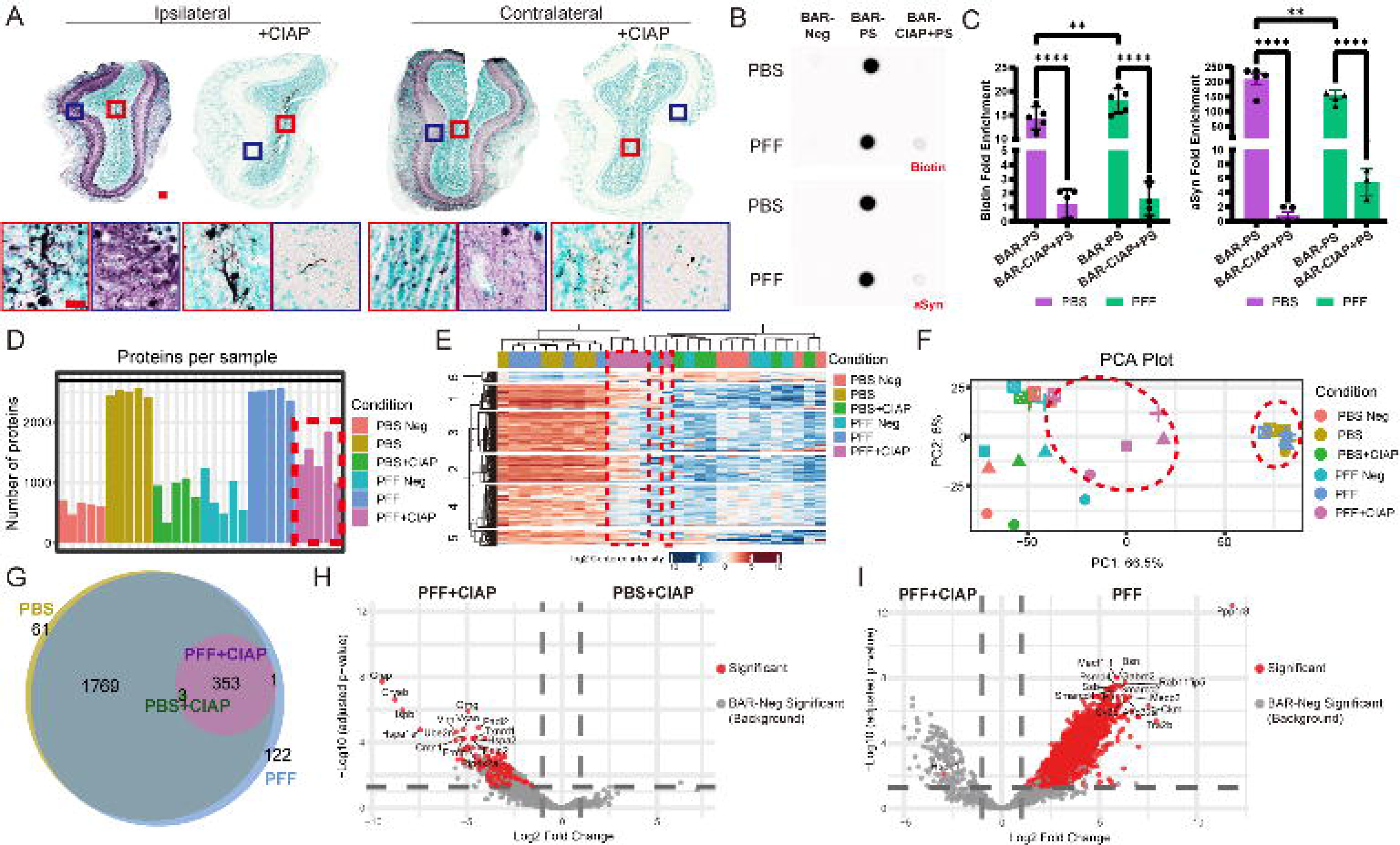
BAR-captured PS129 interactomes in M83 OBs. (A) Representative images of M83 OBs used for BAR capture. PFF-treated OBs from both hemispheres are shown with and without CIAP pretreatment; boxed regions are shown as enlarged 20× views below the corresponding whole-OB images. Scale bars: whole-OB scans, 100 μm; enlarged images, 20 μm. (B) Following BAR labeling, captured proteins were isolated using streptavidin magnetic beads. A fraction of each bead eluate was spotted onto methanol-activated PVDF membranes and probed for biotin and total αsyn. BAR-Neg, primary antibody omission control; BAR-PS, BAR-PS129 capture; BAR-CIAP+PS, BAR capture of CIAP-resistant PS129. (C) Fold enrichment of spot-blot signals for biotin and total αsyn relative to BAR-Neg controls (two-way ANOVA with uncorrected Fisher’s LSD post hoc test; **p < 0.01, ****p < 0.0001, n=5). The remaining bead eluates were analyzed by LC-MS/MS and quantified using MaxQuant LFQ. LFQ Analyst plots show (D) the number of proteins quantified per sample, (E) hierarchical clustering of significant proteins across samples, and (F) PCA of sample-level protein abundance profiles. (G) Interactomes identified in each condition were compared using a weighted Venn diagram. Volcano plots show differential protein abundance for (H) PFF+CIAP versus PBS+CIAP and (I) PFF+CIAP versus PFF, with the top 15 significantly enriched proteins annotated.

**Figure 5.**
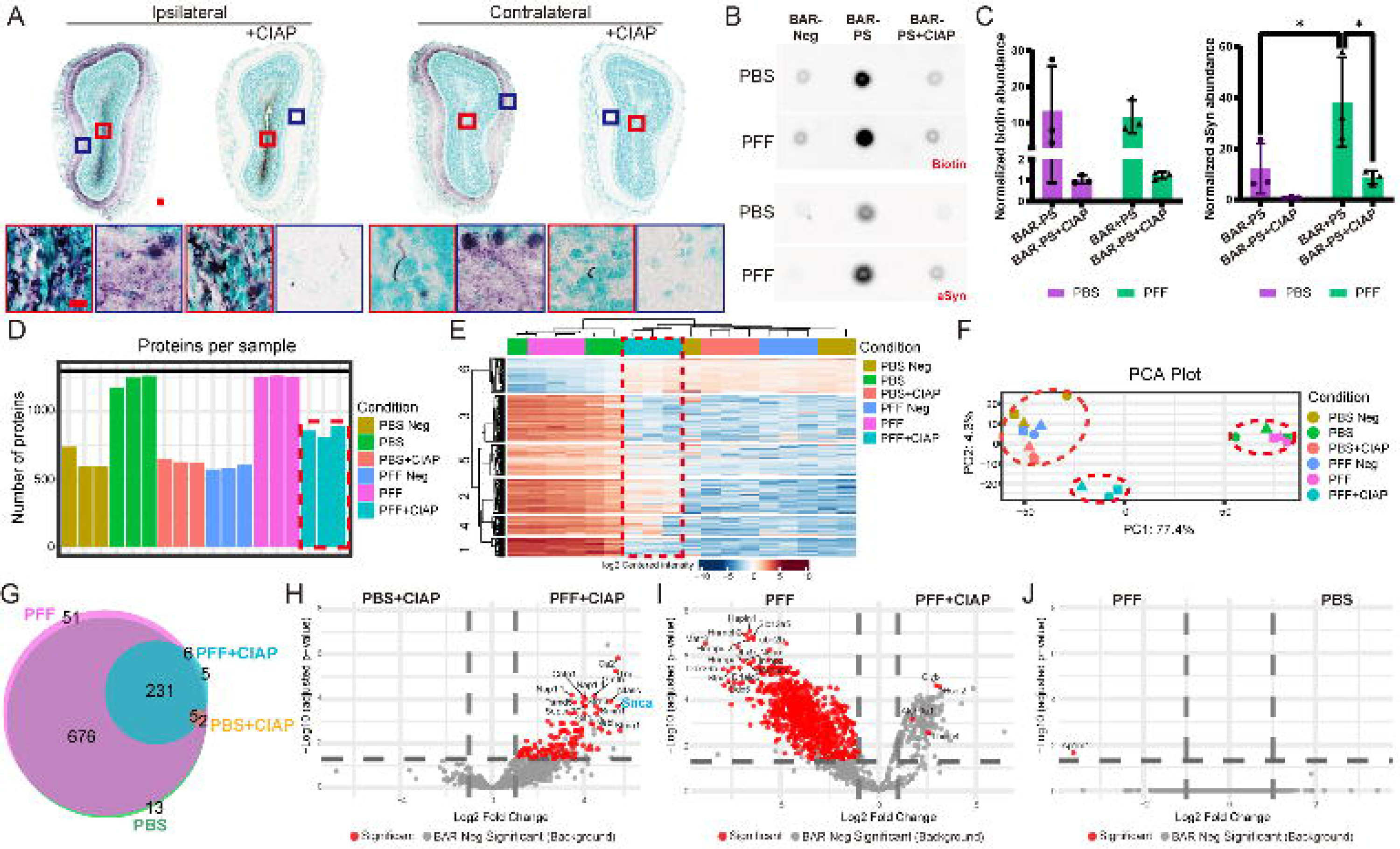
BAR-captured PS129 interactomes in WT OBs. (A) Representative PFF-treated WT OB sections used for BAR capture, shown from both hemispheres with or without CIAP pretreatment. Red and blue boxed regions are shown at 60× magnification. Scale bars: whole-OB scans, 100 μm; magnified images, 10 μm. (B) Spot blot analysis of bead eluate fractions from BAR captures probed for biotin and total αsyn. (C) Quantification of biotin and total αsyn spot blot signals, shown as fold enrichment relative to BAR-Neg controls; two-way ANOVA with uncorrected Fisher’s LSD test, *p < 0.05, n=3. (D) Number of proteins captured per sample. (E) Hierarchical clustering heatmap and (F) principal component analysis plot of all captured samples. (G) Weighted Venn diagram comparing proteins identified across conditions. Volcano plots showing differential protein abundance for (H) PFF+CIAP versus PBS+CIAP, with SNCA highlighted in light blue; (I) PFF+CIAP versus PFF; and (J) PFF versus PBS. The top 15 enriched proteins in each comparison are annotated.

LC-MS/MS detected >2,000 proteins in PBS and PFF groups (Fig. 4D). In contrast, BAR-Neg and PBS+CIAP yielded <750 proteins, while PFF+CIAP samples yielded ∼1,000–2,000 proteins. Hierarchical clustering showed four major clusters.

PFF+CIAP samples clustered well, except for two PFF+CIAP outliers, which clustered closer to the BAR-Neg controls (Fig. 4E; Dataset 1; Fig. S5). PCA revealed tight clustering of PBS and PFF samples along both PC1 and PC2, whereas PFF+CIAP samples exhibited markedly greater dispersion, particularly along PC2 (Fig. 4F)

Differential abundance (DA) relative to BAR-Neg defined the PS129 proximal proteome (Dataset 1; Fig. S6A). Across conditions, 2,309 proteins were identified: 2,186 in PBS, 3 in PBS+CIAP, 2,248 in PFF, and 357 in PFF+CIAP (Fig. 4G; Fig. S6). Most proteins overlapped between conditions (∼92%), with 61 unique to PBS and 123 unique to PFF. PFF+CIAP interactome was largely nested within the PBS and PFF interactomes. Compared with PBS+CIAP, PFF+CIAP was enriched for glial-reactivity, axon–glia/myelin, and proteostatic-stress proteins, including GFAP, CRYAB, HSPB1, HSPA1A, UBE2N, CNTN1, and ERMN (Fig. 4H). HSPB1/Hsp27 was significantly enriched in PFF+CIAP relative to PFF (Fig. 4I). By contrast, DA proteins in the PFF condition were involved in synaptic trafficking and structural organization (BSN, GABRB2, MACF1.1, RAB11FIP5), as well as synaptic homeostasis (CKM, PPP1R8, MECP2, TRA2B, SMARCC2, SSB) (Fig. 4I). Without CIAP, no DA proteins were identified between PFF- and PBS-treated M83 (Fig. S6B; Dataset 1).

### CIAP-resistant PS129 proximal proteome in WT OBs

In PBS-treated OBs, biotin enrichment was 13.25-fold relative to BAR-Neg controls and 1.03-fold after CIAP treatment; in PFF-treated OBs, biotin enrichment was 11.68-fold and 1.23-fold after CIAP treatment (Fig. 5B, C). αSyn enrichment was 12.20-fold in PBS-treated OBs and 38.21-fold in PFF-treated OBs, showing significantly higher enrichment in PFF-treated OBs (two-way ANOVA with uncorrected Fisher’s LSD, p = 0.0136). After CIAP treatment, αsyn enrichment was 0.73-fold in PBS-treated OBs and 8.70-fold in PFF-treated OBs.

LC-MS/MS detected 1,250 proteins in PBS- and PFF-injected WT mice (Fig. 5D). The PFF+CIAP condition yielded ∼750–1,000 proteins, whereas the other conditions yielded ∼500–750 proteins. Hierarchical clustering identified three major clusters, with PFF-CIAP clustered near no CIAP groups (Fig. 5E; dotted line highlights the PFF-CIAP cluster). PCA showed clear condition-dependent separation (Fig. 5F) PBS and PFF samples clustered together, BAR-Neg and PBS+CIAP samples clustered separately, and PFF+CIAP formed a distinct cluster without obvious outliers.

In total, 990 significant interactions were identified across the four conditions: 928 in PBS, 8 in PBS+CIAP, 971 in PFF, and 247 in PFF+CIAP (Fig. 5G, Dataset 2; Fig. S7 for volcano plots BAR-PS129 and BAR-Neg). WT proximal proteome, like those in M83 mice, showed substantial overlap across conditions (∼93% of proteins shared). In contrast, 51 proteins were unique to PFF, and 13 were unique to PBS. PFF+CIAP contained five uniquely identified proteins—ALDH9A1, CLYBL, HSDL2, POLDIP2, and THEM4—all associated with mitochondrial or metabolic function.

DA between the PFF+CIAP and PBS+CIAP conditions revealed enrichment of axon–glia- and extracellular matrix-associated proteins, including NFASC, CNTN1, and TNR, as well as glial-associated proteins such as CA2 and GMFB (5H). Although the enriched proteins differed from M83, both models converged on axon–myelin and glial-associated biology in the aggregate-associated interactome, with SNCA among the top hits.

DA between PFF+CIAP and PFF conditions revealed four mitochondrial or metabolism-associated proteins—CLYBL, HSDL2, ALDH9A1, and THEM4 (Fig. 5I). By contrast, proteins preferentially enriched in the PFF condition were linked to extracellular matrix organization, synaptic ion homeostasis, and nuclear RNA/DNA regulatory processes, including HAPLN1, SLC12A5, HNRNPH2, MATR3, DHX9, NONO, and HNRNPUL2. Only AP1M1 was differentially abundant between PFF and PBS in WT OBs (Fig. 5J), suggesting modest effects on vesicular trafficking or endosomal sorting; no other proteins differed, like M83s.

### Axon–glia-enrichment in PS129 interactomes in M83 mice

Without CIAP, most proteins were shared between PBS and PFF (92%), and 61 and 123 proteins were unique to PBS and PFF, respectively (6A). Shared proteins had similar mean importance scores (IS) values in PBS and PFF conditions (22.35 and 22.21, respectively; Fig. 6B), whereas PBS- and PFF-unique proteins had lower mean IS values (6.65 and 8.18, respectively). Overall, shared proteins had significantly higher IS values than unique proteins (Kruskal–Wallis test followed by Dunn’s multiple comparisons test, ****p < 0.0001).

**Figure 6.**
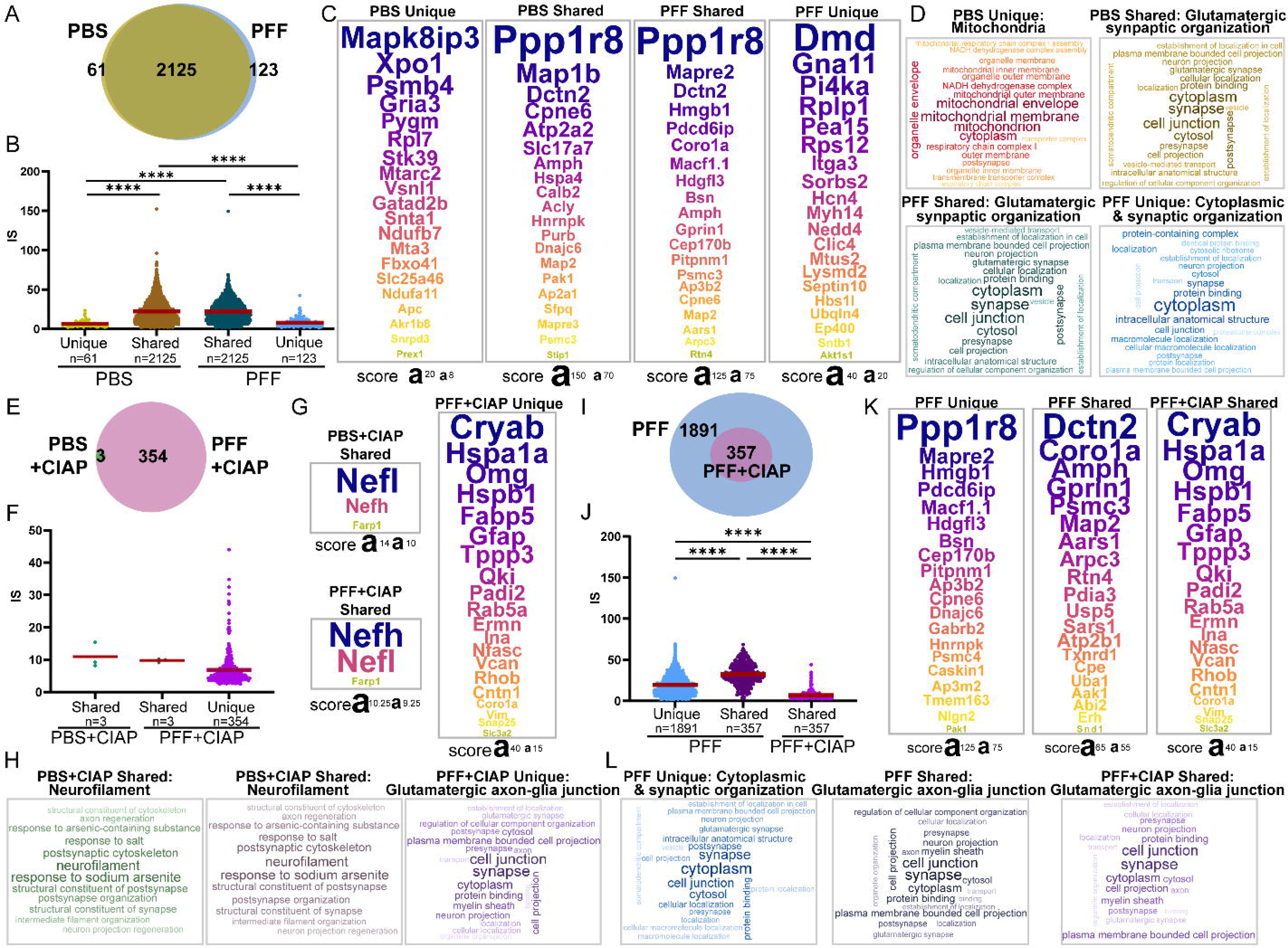
PS129 interactomes in the M83 OB model. (A) Interactomes identified under PBS and PFF conditions were compared using a weighted Venn diagram. A total of 2,125 proteins overlapped between conditions, whereas 61 and 123 proteins were uniquely identified in PBS and PFF, respectively. (B) Importance scores (IS) were compared across PBS-unique, PBS-shared (proteins enriched in PBS and overlapping with the PFF condition), PFF-shared (proteins enriched in PFF and overlapping with the PBS condition), and PFF-unique protein groups. Kruskal–Wallis test with Dunn’s multiple-comparisons test, ****p < 0.0001. (C) Top 20 proteins ranked by IS in each group, with corresponding IS values annotated below. (D) Full ranked protein lists from each group were analyzed in g:Profiler using ordered queries, and the top 20 GO terms are displayed as word clouds. Dominant GO-term themes were manually selected and annotated above each word cloud. (E–H) The same analyses were performed for PBS+CIAP versus PFF+CIAP comparisons and (I–L) for PFF versus PFF+CIAP comparisons. Kruskal–Wallis test with Dunn’s multiple-comparisons test, ****p < 0.0001.

Top 20 proteins ranked by IS revealed MAPK8IP3, an axonal transport scaffold protein linked to PD motor symptoms, as the highest-ranking interaction (Fig. 6C) [53, 54]. Other high-ranking PBS-unique proteins were associated with mitochondrial metabolism (NDUFB7, NDUFA11, SLC25A46, and MTARC2), glutamatergic postsynaptic density organization/signaling (GRIA3, VSNL1, PREX1, SNTA1, and FBXO41), and nuclear-associated processes (XPO1, GATAD2B, MTA3, PSMB4, and STK39). Postsynaptic and nuclear associations are consistent with physiological PS129 interactions in human brain tissue [15].

The PBS-shared group was led by PPP1R8, a nuclear serine/threonine phosphatase PP1 inhibitor, which ranked substantially higher than the next protein (IS: 152.28 vs. 96.12 for MAP1B), reflecting engagement of nuclear regulatory pathways during sustained mutant αsyn overexpression. Other top-ranked PBS-shared proteins were associated with glutamatergic synaptic function (SLC17A7/VGLUT1, CPNE6, AMPH, ATP2A2, AP2A1, DNAJC6, PAK1, MAP2, and CALB2) and cytoskeletal support/transport (MAP1B, DCTN2, and MAPRE3). Notably, DNAJC6/PARK19, a protein required for clathrin-mediated synaptic vesicle recycling and a monogenic cause of early-onset PD, was found in this group [55].

PPP1R8 was also the top-ranked protein in the PFF-shared group. Six of the top 20 proteins overlapped with the PBS-shared group: AMPH, CPNE6, DCTN2, MAP2, PPP1R8, and PSMC3. The top PFF-shared proteins similarly reflected glutamatergic synaptic organization (BSN, AMPH, AP3B2, PITPNM1, CPNE6, MAP2, GPRIN1, and RTN4) and cytoskeletal transport and organization (MAPRE2, DCTN2, MACF1, and ARPC3). Several of these proteins, including BSN, DCTN2, HMGB1, MACF1, and CORO1A, have been implicated in PD- or synucleinopathy-related pathways.

The top 20 PFF-unique proteins identified DMD/dystrophin as the highest-ranking, a protein associated with astrocytic endfeet [56, 57]; other top-ranked proteins, including PEA15, CLIC4, and ITGA3, further support astrocyte-related signaling or adhesion. Consistent with this profile, the PFF-unique group was enriched for proteins associated with axon guidance, glial support, and synaptic stability, including SORBS2, MYH14, ITGA3, SEPTIN10, GNA11, PI4KA, HCN4, CLIC4, AKT1S1, PEA15, and MTUS2, as well as proteotoxic stress-response proteins, including NEDD4, UBQLN4, LYSMD2, EP400, and HBS1L.

GO analysis (Fig. 6D) showed mitochondrial and postsynaptic enrichment in PBS-unique proteins, similar glutamatergic synaptic profiles in shared proteins, and cytoplasmic/synaptic organization enrichment in PFF-unique proteins.

354 proteins identified for aggregate-specific interactome (PFF+CIAP and PBS+CIAP), with only 3 proteins being unique to PBS+CIAP (Fig. 6E). Ranked analysis showed similar mean IS values for the shared PBS+CIAP and PFF+CIAP proteins (10.93 and 9.80, respectively), whereas PFF+CIAP-unique proteins had a mean IS value of 6.82 (Fig. 6F).

The PBS+CIAP and PFF+CIAP groups shared only three proteins: the axonal neurofilaments NEFL and NEFH, and FARP1 (Fig. 6G). In the PFF+CIAP group, two heat shock proteins, CRYAB and HSPB1, were detected. Overall, the top 20 proteins in the CIAP-resistant, aggregate-associated interactome were strongly enriched for glial reactivity and stress-response markers (CRYAB, HSPA1A, HSPB1, GFAP, and VIM), as well as myelin- and axon–glial junction-associated proteins (OMG, QKI, PADI2, ERMN, NFASC, and CNTN1). Top GO terms for PBS+CIAP and PFF+CIAP were neurofilament, while PFF+CIAP-unique proteins were enriched for cell junction, synapse, protein binding, myelin sheath, and cell projection terms, with a dominant theme of glutamatergic axon–glial junctions, consistent with the top 20 proteins (6H).

We then compared PFF+CIAP and PFF conditions. All 357 CIAP-resistant PS129 proteins overlapped with the total PFF interactome (Fig. 6I). IS scores were highest in the PFF-shared group and lowest in the PFF+CIAP-shared group (Fig. 6J; Kruskal–Wallis test followed by Dunn’s multiple comparisons test, ****p < 0.0001). For PFF-unique, PPP1R8 was the top-ranked protein (Fig. 6K). Other high ranked proteins included synapse-associated proteins (BSN, AP3B2/M2, DNAJC6, CPNE6, NLGN2, GABRB2, CASKIN1, and PITPNM1) and cytoskeletal/trafficking proteins (MAPRE2, MACF1, PDCD6IP, and PAK1). For the PFF-shared group, DCTN2, a dynactin complex component, was the top-ranked protein. This group included proteins associated with synaptic organization and vesicle trafficking (AMPH, GPRIN1, AAK1, ABI2, and CPE), cytoskeletal transport/organization (DCTN2, MAP2, ARPC3, and CORO1A), and proteostasis pathways (PSMC3, UBA1, and USP5).

In the PFF+CIAP-shared group, CRYAB was the top-ranked protein, and the top 20 proteins were enriched for glial reactivity and myelin/axon–glia-associated proteins, consistent with Fig. 6G. The GO terms for PFF-unique proteins were enriched for cytoplasmic and synaptic organization terms (Fig. 6L), whereas both the PFF-shared and PFF+CIAP-shared groups were enriched for glutamatergic axon–glial junction-associated terms.

### Axon–glial-enrichment in PS129 interactomes in WT mice

For WT mice, PBS and PFF interactomes largely overlapped, with 914 shared proteins (92.8%) and only 14 and 57 proteins unique to PBS and PFF, respectively (Fig. 7A). The PFF-shared group had a significantly higher mean IS than the PBS-shared group (21.37 vs. 18.16; Fig. 7B; Kruskal–Wallis test followed by Dunn’s multiple comparisons test, **p < 0.0001), reflecting both higher overall abundance and stronger statistical support for the same proteins in PFF (Dataset 2). PBS- and PFF-unique proteins had lower mean IS values of 7.40 and 8.92, respectively. Top-ranked proteins were HBB-B2/hemoglobin beta-2 for PBS-unique, Matr3 for PBS/PFF shared, and Cap1 for PFF unique, as the highest-ranking protein (Fig. 7C). PBS-unique group proteins associated with synaptic/postsynaptic organization (ACTN2, PTPRS, ABI2, AFDN, PPM1E, and BCAN), along with axonal cytoskeletal, nuclear, metabolic, and glial-associated proteins (NEFM, LMNB1, DYNLL2, S100A5, PYGM, and FABP7).

**Figure 7.**
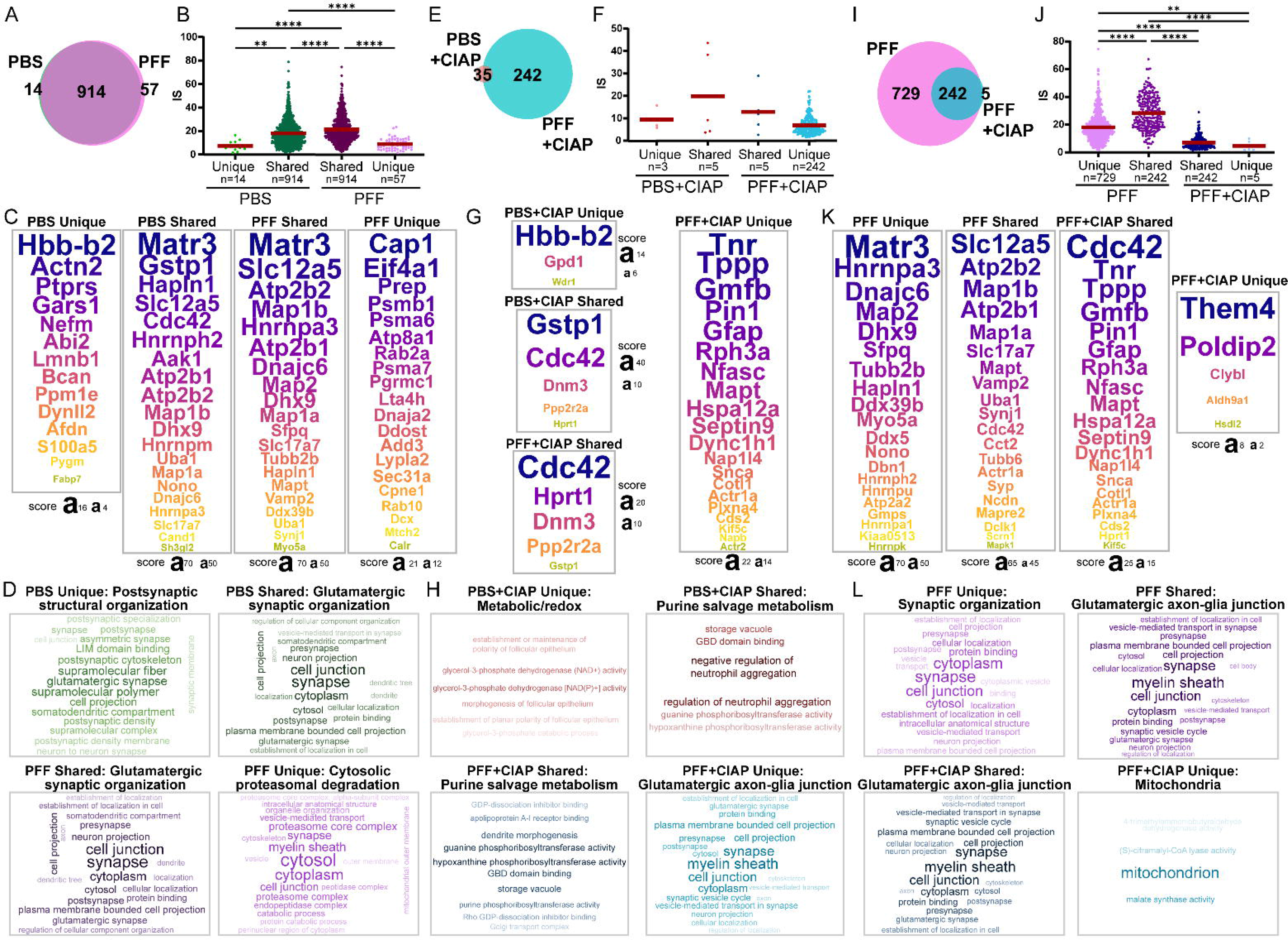
PS129 interactomes in the WT OB model. (A) PBS and PFF interactomes were compared using a weighted Venn diagram. A total of 985 proteins were identified, with 914 overlapping between conditions, whereas 14 and 57 proteins were uniquely identified in PBS and PFF, respectively. (B) IS values were compared across PBS-unique, PBS-shared, PFF-shared, and PFF-unique protein groups. Kruskal–Wallis test with Dunn’s multiple-comparisons test, **p < 0.01, ****p < 0.0001. (C) Top 20 proteins ranked by IS in each group are shown, with corresponding IS values annotated below. (D) The top 20 GO terms from the full ranked protein lists for each group are displayed as word clouds. Overarching GO-term themes were manually selected and annotated above each word cloud. (E–H) The same analyses were performed for PBS+CIAP versus PFF+CIAP comparisons. (I–L) The same analyses were performed for PFF versus PFF+CIAP comparisons. Kruskal–Wallis test with Dunn’s multiple-comparisons test, **p < 0.01, ****p < 0.0001.

MATR3 was the top-ranked protein in the PBS-shared and PFF-shared group (Fig. 7C). PBS-Shared was enriched for glutamatergic synaptic proteins, including SLC17A7, SH3GL2, DNAJC6, AAK1, CDC42, MAP1A, MAP1B, ATP2B1, ATP2B2, and HAPLN1, along with RNA/nuclear regulatory and proteostasis-related proteins, including HNRNPH2, DHX9, HNRNPM, NONO, HNRNPA3, UBA1, and CAND1. The PFF-shared group was also enriched for glutamatergic synaptic and RNA/nuclear regulatory proteins, as well as a neuronal projection/cytoskeletal signature, including microtubule-associated proteins (MAP1B, MAP2, MAP1A, and MAPT) and TUBB2B. This suggests that PFF seeding preferentially engages axonal- and cytoskeleton-associated proteins in WT OBs. CAP1, the top-ranked PFF-unique protein, suggests cytoskeletal remodeling, while the remaining hits point to proteostasis and intracellular trafficking consistent with neuronal responses to PFF-seeded pathology. There was strong enrichment for glutamatergic synapses in the PBS-unique group (Fig. 7D). In contrast, the PFF-unique group was enriched for cytosol, cytoplasm, myelin sheath, cell junction, and proteasome core complex terms, suggesting cytosolic proteasome-mediated proteostasis with synaptic/cytoskeletal and myelin-associated organization.

Next, we compared CIAP-resistant PS129 conditions. PBS+CIAP contained eight identified proteins, five of which overlapped with PFF+CIAP, whereas 242 proteins were unique to PFF+CIAP (Fig. 7E). Mean IS values were highest for PBS+CIAP-shared proteins (19.84), followed by PFF+CIAP-shared (12.86), PBS+CIAP-unique (9.44), and PFF+CIAP-unique proteins (6.86; Fig. 7F). The PBS+CIAP-unique proteins—HBB-B2, GPD1, and WDR1—suggested blood/vascular signal, metabolic regulation, and actin cytoskeletal remodeling, respectively (Fig. 7G). Shared PBS+CIAP/PFF+CIAP proteins included GSTP1, CDC42, DNM3, PPP2R2A, and HPRT1, reflecting oxidative stress, cytoskeletal/synaptic regulation, vesicle trafficking, phosphatase signaling, and purine metabolism. In the aggregate-specific PFF+CIAP-unique interactome, TNR, an oligodendrocyte-enriched extracellular matrix glycoprotein expressed during myelination, was the top-ranked protein [58, 59]. This group was enriched for axon–glia/glial-reactivity proteins (TNR, NFASC, GFAP, GMFB, and TPPP) and synaptic trafficking, cytoskeletal, and axonal transport proteins, including SNCA/αsyn, the BAR target, along with RPH3A, NAPB, MAPT, PIN1, DYNC1H1, KIF5C, ACTR1A, and ACTR2. Although the biological themes resembled the M83 PFF+CIAP interactome (Fig. 6G), only GFAP and NFASC overlapped among the top 20 proteins between models.

For GO analysis, the PBS+CIAP group was dominated by metabolism and redox balance terms (Fig. 7H). PBS+CIAP- and PFF+CIAP-shared proteins were enriched for purine salvage metabolism, driven mainly by hypoxanthine- and guanine-associated terms. In contrast, PFF+CIAP-unique proteins were enriched for myelin sheath, cell junction, synapse, and cell projection terms, suggesting glutamatergic axon–glia junction-associated pathways.

Lastly, we compared the PFF+CIAP interactome with the total PFF interactome (Fig. 7I). Of the PFF+CIAP proteins, 242 overlapped with the PFF interactome, whereas five were unique to PFF+CIAP and 729 were unique to PFF. The PFF-shared group had the highest mean IS value (28.32; Fig. 7J), followed by the PFF-unique group (18.08), while the PFF+CIAP-shared and PFF+CIAP-unique groups had lower mean IS values of 7.03 and 4.69, respectively (Kruskal–Wallis test followed by Dunn’s multiple comparisons test, **p < 0.01, ****p < 0.0001).

Top-ranked PFF-unique proteins included MATR3 (Fig. 7K), and this group was enriched for RNA-binding/nuclear regulatory proteins (MATR3, HNRNPA3, DHX9, SFPQ, DDX39B, DDX5, NONO, HNRNPH2, HNRNPU, HNRNPA1, HNRNPK) and neuronal cytoskeletal/synaptic structure proteins (MAP2, TUBB2B, DBN1, MYO5A, DNAJC6, HAPLN1, ATP2A2).

In the PFF-shared group, SLC12A5/KCC2, a regulator of neuronal chloride homeostasis and excitability, was the highest-ranking protein. This group was enriched for synaptic vesicle/glutamatergic synaptic proteins (SLC17A7/VGLUT1, VAMP2, SYP, SYNJ1), cytoskeletal/axonal organization proteins (MAP1B, MAP1A, MAPT, MAPRE2, TUBB6, DCLK1, ACTR1A), and calcium/ion homeostasis proteins (ATP2B1, ATP2B2, SLC12A5).

In the PFF+CIAP-shared group, only ACTR1A, CDC42, and MAPT overlapped with the PFF-shared top 20. CDC42, the top-ranked protein, is a cytoskeletal/signaling Rho-family GTPase reported to interact with αsyn and to be pharmacologically targetable in PD animal models [60, 61]. The top 20 proteins also included SNCA/αsyn, along with axon–glia/myelin- and glial-associated proteins (NFASC, TNR, TPPP, GFAP, GMFB) and axonal transport/cytoskeletal proteins (DYNC1H1, ACTR1A, KIF5C, MAPT, PLXNA4).

When PFF+CIAP-shared proteins were cross-ranked against the PFF-shared group, there IS ranks ranged from MAPT at rank 7 to CDS2 at rank 112. Within the broader PFF-enriched set, they remained relatively high-ranking, from MAPT at rank 15 to CDC42 at rank 218. In contrast, their ranks in the total PBS-enriched set were more variable, from CDC42 at rank 5 to GFAP at rank 753 (Dataset 3, Fig. S8B). Similar to the M83 model, these patterns suggest that most of the aggregate-associated interactome reflects selective prioritization within the physiological PS129 interactome; in WT OBs, this prioritization is already partially reflected in the total PFF-enriched PS129 interactome.

Among the five proteins uniquely identified in the PFF+CIAP group, THEM4, a mitochondria-associated signaling regulator, was the top-ranked protein. The remaining proteins—CLYBL, ALDH9A1, HSDL2, and POLDIP2—were also mitochondria-associated or metabolic proteins. Although small, this newly detected WT PFF+CIAP-unique subset suggests a pathology-linked mitochondrial/metabolic component.

GO analysis showed that the PFF-unique group was enriched for synapse, cell junction, cytoplasm, protein binding, and neuron projection terms, consistent with broad synaptic organization (Fig. 7L). Proteins shared between PFF and PFF+CIAP were enriched for myelin sheath, cell junction, synapse, cytoplasm, and glutamatergic synapse terms, suggesting involvement in glutamatergic axon–glial junction-related processes. In contrast, the PFF+CIAP-unique group was dominated by mitochondria-associated pathways.

### Axon–glial pathways between two synucleinopathy mouse models

For mice treated with PBS, 40% of proteins were shared between models (Fig. 8A). Only 34 proteins were unique to WT, whereas 1,292 were unique to M83. Shared proteins showed a moderate positive correlation in IS between models (r ≈ 0.333, R² = 0.111, p < 0.001; Fig. S9A). A similar pattern was observed under PFF-treated conditions (Fig. 8B), where shared proteins accounted for 41% of all PFF-enriched proteins across both models; only 33 proteins were unique to WT, whereas 250 were unique to M83. IS values among shared proteins were again moderately correlated between models (r ≈ 0.346, R² = 0.120, p < 0.001; Fig. S9B).

**Figure 8.**
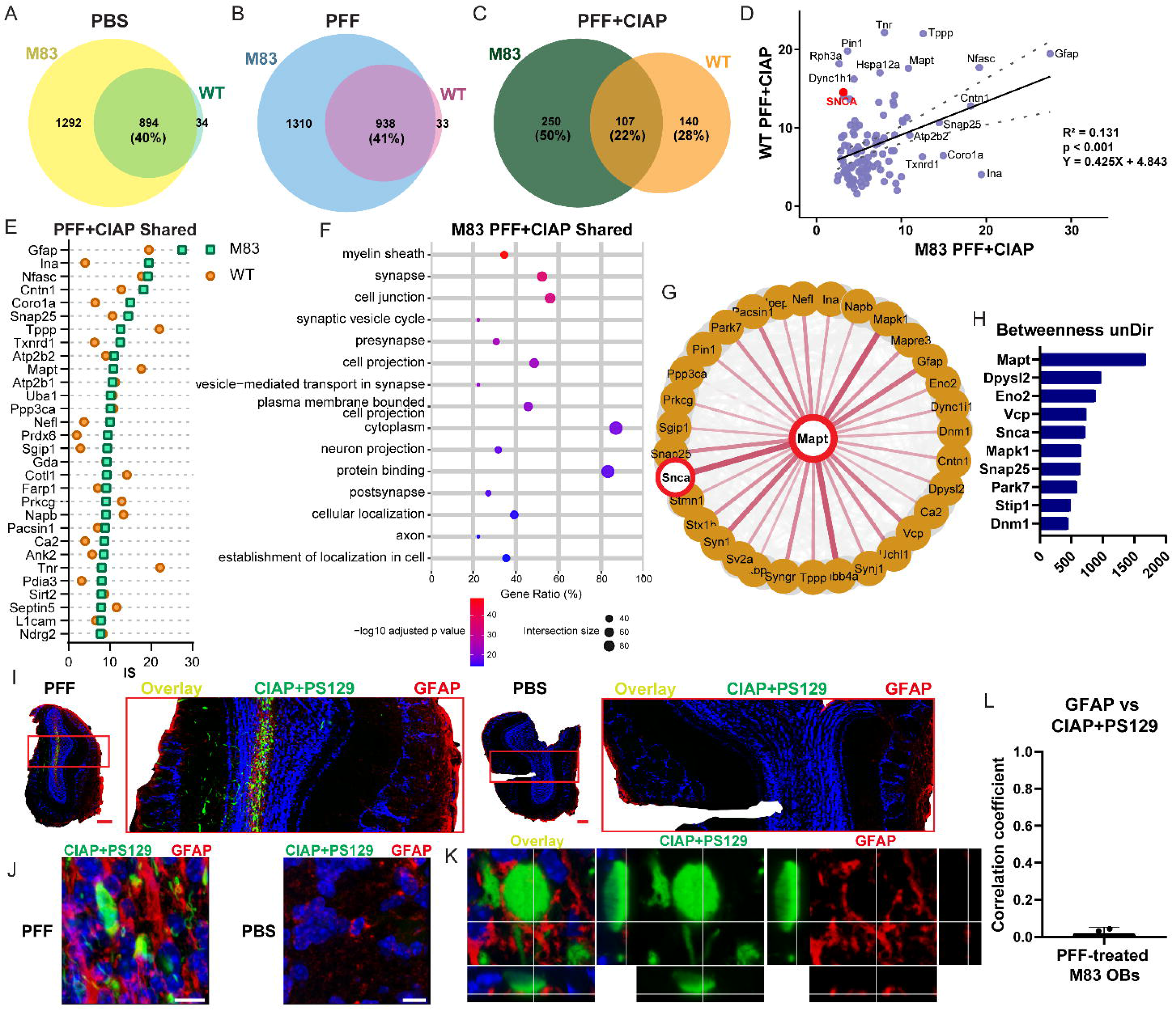
Consensus pathway for PFF-seeded αsyn interactomes. (A–C) Weighted Venn diagrams comparing enriched proteins identified under (A) PBS, (B) PFF, and (C) PFF+CIAP conditions in M83 and WT mice. (D) Correlation plot comparing IS values for the 107 PFF+CIAP-shared proteins between models. (E) Top 30 M83 PFF+CIAP-shared proteins ranked by IS, with corresponding WT IS values shown. (F) Top 15 GO terms for the 107 shared proteins, analyzed in g:Profiler using M83 IS-ranked proteins and ordered by significance. (G) Cytoscape STRING functional interaction network of the 107 shared proteins, followed by CentiScaPe 2.2 centrality analysis, identified MAPT as the central hub protein based on betweenness unDir score. The STRING confidence score cutoff was set to 0.4. Nodes represent proteins, and edges represent interaction confidence scores. First neighbors of MAPT based on undirected interactions are labeled, while all other interacting proteins and interactions are shown in light gray. (H) The top 10 proteins ranked by betweenness unDir score among the consensus proteins are shown. (I) Multiplex labeling of GFAP and CIAP-resistant PS129 in PFF- and PBS-treated M83 OBs. Red-boxed regions are enlarged next to whole-OB images. Scale bars = 200 μm. (J) Representative 60× z-stacked maximum-intensity projections of the OB GCL showing dysmorphic CIAP-resistant PS129-positive neurites and closely apposed GFAP-positive astrocytes. Scale bars = 10 μm. (K) Slice view showing GFAP-positive signals surrounding Lewy body-like structures, with signal distribution shown in the x, y, and z planes. (L) Pearson’s correlation coefficient for GFAP versus CIAP-resistant PS129 signals in M83 OBs. Mean = 0.018, SD = 0.034, n = 3.

In contrast, the PFF+CIAP condition showed substantially less overlap between models: only 22% of proteins were shared, whereas 50% and 28% were unique to M83 and WT, respectively (Fig. 8C). Among the 107 shared proteins, IS values were moderately correlated between models (r ≈ 0.362, R² = 0.131, p < 0.001; Fig. 8D). Within this shared PFF+CIAP set, GFAP, a reactive astrocyte marker, had the highest IS in M83 OBs, whereas TNR, an oligodendrocyte-enriched extracellular matrix protein, was top-ranked in WT OBs. Overall, both models showed prominent glial and axon–glia-associated proteins, including TPPP, MAPT, CNTN1, CORO1A, and NFASC. SNCA also ranked among the top 10 proteins by IS in WT OBs (Fig. 8D, red dot).

The 107 consensus CIAP-resistant aggregate-specific proteins were examined further. The top 30 consensus PFF+CIAP proteins in M83 OBs were compared with their corresponding WT IS values (Fig. 8E; full list in Dataset 3). GFAP, a marker of reactive astrocytes, had the highest IS in M83 OBs (27.52). The shared interactome included axon–glia/adhesion proteins (NFASC, CNTN1, TNR, L1CAM), glial-reactivity proteins (GFAP, CORO1A, VIM), axonal/cytoskeletal proteins (INA, NEFL, MAPT, ANK2, SEPTIN5), oligodendrocyte/myelin-associated proteins (TPPP, CA2), and synaptic proteins involved in glutamatergic signaling and vesicle biology (SNAP25, FARP1, NAPB, PACSIN1), consistent with the role of synaptic vesicle pathways in αsyn biology [62–64]. Although these proteins were shared between the M83 and WT PFF+CIAP profiles, their relative IS rankings differed (Fig. 8E), indicating model-specific prioritization within a shared aggregate-associated interactome. Overall, these findings further support an association between CIAP-resistant aggregates and axon–glia/myelin-associated pathways, as well as glutamatergic synaptic biology.

GO enrichment analysis of the full ranked set of 107 consensus M83 proteins further supported these themes (Fig. 8F). The leading GO terms included myelin sheath, synapse, cell junction, synaptic vesicle cycle, neuron projection, and axon. The same shared protein set, ranked by WT IS values, showed highly similar GO enrichment, with only minor shifts in selected term order (Fig. S9C).

The consensus aggregate-specific interactome was analyzed using Cytoscape and the STRING database, and network centrality metrics. This analysis identified a primary cluster of 95 proteins (Fig. 8G; complete functional interactome shown in Fig. S10). MAPT/Tau showed the highest undirected betweenness centrality score (1,980). This finding is consistent with the BAR-PS129 interactome identified in PD/DLB brains [28]. Several additional high-centrality proteins also have established links to PD or related neurodegenerative pathways, including VCP, an ATPase involved in proteostasis and misfolded-protein clearance linked to Parkinsonism [65]; SNCA/αsyn; PARK7/DJ-1, an oxidative-stress and mitochondrial-protective protein associated with early-onset familial PD [66]; and SNAP25, DPYSL2, and MAPK1, which have been implicated in synaptic, axonal, or neurodegenerative processes relevant to PD/DLB [67, 68]. Together, these findings suggest that the shared CIAP-resistant PS129 interactome in the OB is organized around axonal structure, proteostasis, and cellular stress-response pathways.

Because GFAP was the top-ranked protein in the consensus aggregate-specific interactome in M83 OBs, we performed multiplex labeling for GFAP and CIAP-resistant PS129. In PFF-treated OBs, both GFAP and CIAP-resistant PS129 signals were concentrated in the GCL, corresponding to the injection site, whereas PBS-treated OBs showed no detectable CIAP-resistant PS129 and substantially lower GFAP labeling (Fig. 8I). Higher-magnification z-stack maximum-intensity projections confirmed abundant GFAP-positive astrocytic processes closely opposed to dysmorphic CIAP-resistant PS129 pathology in PFF-treated OBs, while PBS-treated OBs showed minimal GFAP labeling (Fig. 8J). Single-plane orthogonal views further showed GFAP-positive processes positioned around CIAP-resistant PS129-positive Lewy body-like structures, without clear signal overlap. Consistent with these observations, Pearson’s correlation analysis showed minimal colocalization between GFAP and CIAP-resistant PS129 signals in OBs (Fig. 8L; mean Pearson’s correlation coefficient = 0.018, SD = 0.034, n = 3). Together, these findings indicate that although GFAP-positive astrocytic processes do not directly colocalize with CIAP-resistant PS129 pathology, robust astrocytic reactivity is spatially concentrated in regions containing CIAP-resistant PS129-positive aggregates in PFF-treated M83 OBs.

## Discussion

Using proximal proteome, we systematically characterize the molecular environment of PS129 as it transitions from a physiological to an aggregated state in two OB-PFF models. Prior studies linked physiological αsyn and PS129 to a wide range of interactions, including components of endocytosis and vesicle trafficking, RNA metabolism, SNARE proteins, cytoskeleton, and mitochondria [13, 15, 69, 70], with aggregate-associated interactomes largely overlapping physiological networks [27, 28, 71, 72]. We identified state-specific PS129-proximal proteomes, including a 107-protein CIAP-resistant aggregate signature enriched for myelinated axon-glial interfaces and retrograde axonal transport motors, consistent with OB glutamatergic centrifugal axons acting as primary seeding/spread conduit, as summarized in Figure 9. Because the olfactory bulb and tract are affected early (Braak I), including in prodromal synucleinopathies (e.g., incidental Lewy pathology), these mechanisms provide molecular context for early αsyn aggregation. While we focused on top-ranked proteins, the hundreds of significant state-specific PS129 interactions identified here offer a valuable resource for future studies.

**Figure 9.**
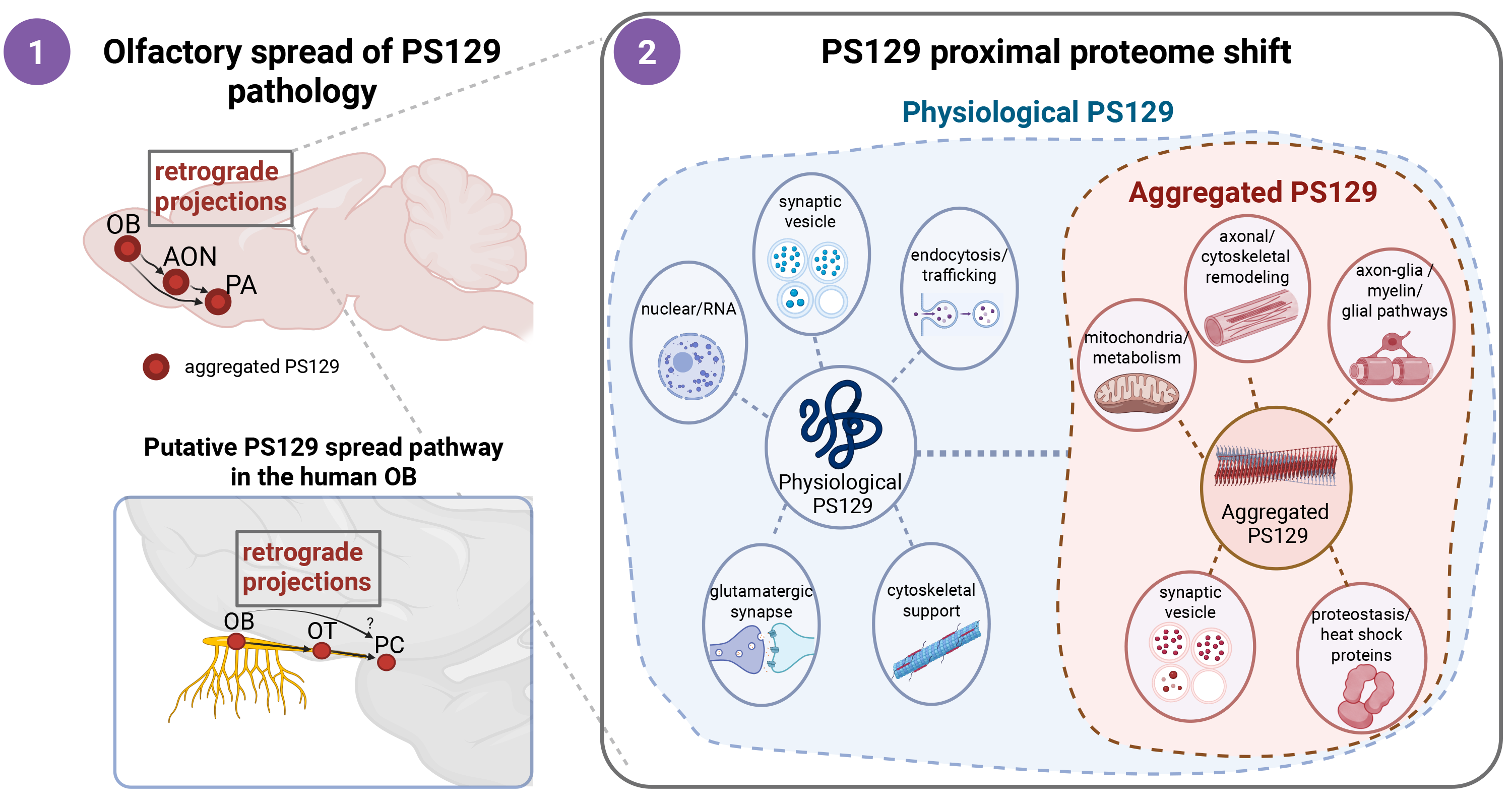
Olfactory circuit-associated spread and disease-associated remodeling of the PS129 proximal proteome. Schematic summarizing the proposed model. Aggregated PS129 pathology may spread through olfactory-connected centrifugal circuits. In mouse models, PS129 pathology propagates retrogradely along projections from the PA and AON to the OB GCL. In human disease, aggregated PS129 is detected in the OB and olfactory peduncle (OP), which contains AON subdivisions, suggesting possible involvement of retrograde OB–AON circuit connections. Pathology within the AON may then extend retrogradely to additional olfactory-connected regions, including the piriform cortex (PC), but whether the human PC provides direct centrifugal input to the OB remains unclear. Right, proximity proteomics suggests that aggregated PS129 is not built from a separate molecular network, but instead reflects disease-associated remodeling of the broader physiological PS129 proximal proteome. Physiological PS129 is associated with synaptic vesicle biology, including SNAREs, endocytosis/trafficking, nuclear/RNA-related proteins, glutamatergic synapse pathways, and cytoskeletal support, whereas aggregated PS129 is enriched for axonal/cytoskeletal remodeling, axon–glia and myelin-associated biology, mitochondrial metabolism, synaptic vesicle pathways, and proteostasis/heat shock responses. Together, this model suggests that pathological PS129 represents a redirected subset of physiological PS129 interactions within vulnerable olfactory circuits. Schematic created in BioRender. Created in BioRender. Choi, S. (2026) https://BioRender.com/3q8nqq5

### Seeding, aggregation, and spread in OB-PFF models

isSID-reactive species closely tracked CIAP-resistant PS129 pathology in M83 mice, whereas WT mice showed limited CIAP-resistant pathology and minimal isSID signal (Figs. 3, S3, and S4). However, isSID signal was not uniformly distributed across CIAP-resistant pathology. Instead, it appeared as discrete puncta or granular foci within, around, or along PS129-positive neurites (Fig. 3B–F), indicating only a small subset of aggregates at discrete foci are actively seeding, and that this seeding is spatially distinct from gross αsyn accumulation (i.e., CIAP-resistant PS129), consistent with prior reports of limited isSID–PS129 overlap [73].

This dissociation aligns with evidence that short or fragmented αSyn fibrils exhibit higher seeding activity, whereas maturation reduces fragmentation and seeding potential [74–77]. Thus, isSID likely detects a dynamic, seed-competent pool of misfolded αSyn associated with, but distinct from, mature CIAP-resistant aggregates. As aggregates mature, seed-competent species become sequestered into stable assemblies with reduced propagation capacity [74–77]. Nevertheless, mature aggregates may still contribute to neurodegeneration by disrupting mitochondria, membranes, and synapses, while also serving as a reservoir for smaller soluble toxic αSyn oligomers generated through dynamic surface and edge dissociation, particularly upon neuronal membrane interaction [23, 77, 78].

### Seeding and aggregation shifts the PS129 proximal proteome

Most PS129 interactions overlapped BAR conditions for both animal models, suggesting that PS129-aggregate interactions are predominantly a reprioritization of canonical PS129 interactions. These aggregate-associated interactions were enriched for axonal/cytoskeletal remodeling, glial support, myelin-associated organization, and proteasome pathways, suggesting an adaptive response to seeded axonal pathology.

In M83 mice, PFF-unique proteins included NEDD4, an E3 ubiquitin ligase that promotes αsyn degradation, and WDR44/Rab11BP, a recently identified modulator of early aSyn aggregation at the lysosomal membrane [79]. The broader M83 PFF interactome also contained PPM1H (higher IS than in PBS) and multiple Rab GTPases, supporting the involvement of the LRRK2–RAB–PPM1H axis in axonal autophagosome transport and αSyn degradation [80]. WT PFF-unique proteins included PREP/prolyl oligopeptidase, which promotes αsyn dimerization/aggregation, and RAB10, a major LRRK2 substrate linked to extracellular αsyn aggregate release [81–83]. Notably, PREP and RAB10 were also present in M83 PBS and PFF interactomes, with PREP potentially resulting from oligomer [84] or A53T αsyn interactions in M83 mice. The M83 aggregated PS129 interactome contained 357 proteins, 356 of which overlapped with the PBS interactome (Fig. S8A), with top-ranked hits dominated by heat shock and glia-associated proteins. Among the top five were CRYAB/HSPB5, HSPA1A/HSP70, and HSPB1/HSP27 (Fig. 6G). HSPB1, repeatedly proposed as a therapeutic candidate for pathological αsyn aggregation and limits fibril elongation/cytotoxicity [85–87].

CRYAB/HSPB5, a chaperone detected in Lewy pathology in PD and abundant in glial cytoplasmic inclusions in MSA, limits protein misfolding and aggregation, can promote HSP70-mediated αsyn disaggregation, and is generally interpreted as part of a protective response to αsyn fibril formation [85, 88–91]. Together, these HSPs suggest engagement of proteostatic pathways that may counteract pathological αsyn aggregation.

In contrast, the WT aggregated PS129 interactome contained 247 proteins, 236 of which overlapped with PBS, leaving 11 PBS-absent proteins (Fig. S8B). Five of these were also absent from PFF and therefore unique to aggregated PS129 (Fig. 7I); all were mitochondrial or metabolic. However, this likely reflects the exacerbation of pre-existing mitochondrial associations rather than the recruitment of a new functional class, as the WT PBS interactome already included MTND4, TOMM70, VDAC1, MTX2, CISD1, RHOT1, DNM1L, SNPH, and PARK7/DJ-1 [92]. The remaining six PBS-absent proteins overlapped with PFF and mapped to mitochondrial, neurofilament, myelin, and ubiquitin-associated functions, consistent with the broader WT aggregated PS129 interactome.

### PS129 at the axon-glial interface

Although the models showed distinct PS129 pathology distributions, they shared a conserved 107-protein aggregated PS129 interactome primarily enriched for myelin sheath, cell-junction, and axonal terms (Figs. 8F, G, and S9C). MAPT/Tau, biochemically and pathologically linked to αsyn and Lewy pathology, had the highest betweenness centrality, suggesting that conserved aggregated PS129 networks center on axonal pathology and the axon–glia interface [93–95]. All 107 consensus proteins appeared in M83 PBS and 103 in WT PBS, further indicating that seeded pathology preferentially engages axonal–glial subsets of physiological PS129 networks.

GFAP emerged as the top consensus aggregate-specific protein in M83 OBs; we did not find evidence of PS129-positive astrocytes (Fig. 8L). Given that BAR captures proteins in proximity [26, 96, 97], GFAP enrichment likely reflects abundant astrocytic processes closely opposed to aggregated PS129 pathology rather than direct colocalization. This is further supported by concurrent enrichment of HEPACAM (astrocytic end foot protein), and NRCAM (neuron–astrocyte adhesion molecule) in the aggregated PS129 interactome [98, 99]. Astrocytes did not have CIAP-resistant PS129, consistent with astrocytes internalizing PFFs but not developing αsyn aggregates [100–103]. High GFAP enrichment observed likely originates from the glial-axon interface, which can be very close (∼10 nanometers) [104], well within the BAR labeling radius.

PFF exposure can induce local pathology in OB granule cells through uptake [10], but the strong GCL signal does not fully explain the consistent enrichment of glutamatergic synapse-associated terms in the aggregated PS129 interactome (Figs. 7L and 8L), given that the GCL is dominated by axonless inhibitory granule cells [105, 106]. Instead, this signature likely reflects afferent or centrifugal projection fibers within the GCL. Consistent with this, OB PFF injection shows that by ∼3 months, pathology propagates mainly along retrograde centrifugal inputs targeting granule cells [6]. In our model, aggregated PS129 was prominent in the AON and APC (Fig. 3A), regions reciprocally connected with the OB through glutamatergic olfactory circuits [9, 10].

This interpretation aligns with preferential retrograde spread of PFF-induced αsyn pathology, with ∼4–4.5-fold greater retrograde than anterograde [107], likely reflecting more efficient dynein-mediated transport of endocytosed PFF cargo toward the soma, than kinesin-mediated anterograde transport [108]. Consistently, DYNC1H1 and DYNC1I1 were detected across both models, with broader dynein representation in M83, whereas kinesin detection was more limited, with KIF21A identified in WT and KIF5C in M83. This pattern may be relevant to human synucleinopathy, in which the OB and AON are among the earliest sites of aggregated PS129 [3, 109–112]. In human PD/DLB OBs, CIAP-resistant aggregated PS129 was strongly enriched in the central OB, particularly the bulbar AON (AONb; Fig. S11) [110, 111, 113]. Olfactory peduncle (OP) sections containing rostrocaudal AON subdivisions also showed abundant PS129 pathology, reinforcing early OB–AON circuit involvement [110, 111, 113]. As in mouse models, these human regions contained seed-competent aSyn species (Fig. S12), indicating that olfactory structures harbor propagation-competent substrates. Because the human AON and piriform cortex are dominated by pyramidal glutamatergic neurons [9, 10], our findings suggest that OB PS129 aggregates may spread retrogradely through glutamatergic synapses within connected olfactory pathways.

This vulnerability is further supported by the enrichment of αSyn at excitatory synapses and its role in modulating seeded aggregation, suggesting that interconnected olfactory pathways may be particularly susceptible to circuit-based propagation of αSyn pathology [114–116]. Consistent with this possibility, the aggregated PS129 proximal proteome was enriched for myelin-, glial-, and proteostasis-associated proteins, including CNP, TPPP, UBA1, and UCHL1. UBA1 and UCHL1 implicate ubiquitin-associated proteostasis within the axon-enriched aggregated PS129 network [92]. More broadly, selective inhibition of neuron-specific plasma membrane-bound neuroproteasomes has been shown to promote endogenous tau aggregation, suggesting that local membrane-associated protein quality-control systems can influence pathological protein aggregation; however, whether analogous mechanisms operate in αSyn pathology remains unclear.

### Limitations

CIAP pretreatment may exclude intermediate, less-phosphatase-resistant αsyn aggregate species, thus underestimating aggregates. Second, model-specific factors may also influence interpretation. For example, although M83 mice are prone to αsyn aggregation, the ectopic expression of A53T αsyn, under the prion promoter, likely results in false positives, with unclear biological relevance to human synucleinopathy. WT mice develop less αsyn pathology but are the closest approximation of the physiological PS129 interactome. To mitigate this model-specific bias, we focus on BAR-identified protein interactions shared between M83 and WT mice.

### Conclusion

Collectively, our results demonstrate that αsyn aggregation largely engages a subset of the non-aggregated PS129 network enriched at the axon–glial interface. This may reflect an active glial response to OB aggregate-bearing neurites of glutamatergic centrifugal circuits, mirroring early OB/AON involvement in human synucleinopathy.

These insights delineate distinct molecular phases of synucleinopathy progression and identify axon–glia interfaces as key therapeutic targets.

## Acknowledgements

This research was supported by the National Institutes of Health under award number 5R01NS128467 and 1RF1NS146420-01 as well as through generous philanthropic support.

