## Supplemental for "Preformed Fibril Seeding Reshapes the Phosphorylated α-Synuclein Proximal Proteome in the Olfactory Bulb"


**This PDF file includes:**


**Figure S1**. Figure S1. RGB thresholding workflow for PS129 percent-area quantification
**Figure S2**. Additional characterization of CIAP-resistant PS129 pathology in the WT-PFF model.
**Figure S3**. Characterization of iSAA signal in PFF-seeded mouse models.
**Figure S4**. Multiplex tyramide signal amplification labeling of iSAA-reactive species and CIAP-resistant or total PS129 pathology.
**Figure S5**. CIAP-resistant PS129 labeling in OB sections used for BAR analysis.
**Figure S6.** Volcano plot analysis of BAR-enriched proteins in M83 OB samples.
**Figure S7**. Volcano plot analysis of BAR-enriched proteins in WT OB samples.
**Figure S8**. Comparison of PBS and PFF+CIAP PS129 interactomes in M83 and WT mice.
**Figure S9**. Model-specific and shared features of WT and M83 PS129 interactomes.
**Figure S10**. Functional interaction network of the 107 consensus CIAP-resistant PS129-associated proteins.
**Figure S11**. CIAP-resistant PS129 pathology in PD/DLB OB and OP.
**Figure S12**. iSAA detection in PD/DLB OB and OP
**Table S1**. Combined wet mass of M83 and WT OB samples used for BAR captures.


**Other supporting materials for this manuscript include the following:**


**Dataset S1 (separate file).** LFQ-Analyst full proteome output from M83 OB BAR samples.
**Dataset S2 (separate file).** LFQ-Analyst full proteome output from WT OB BAR samples.
**Dataset S3 (separate file).** Importance score-ranked BAR-enriched proteins from M83 and WT OB datasets.

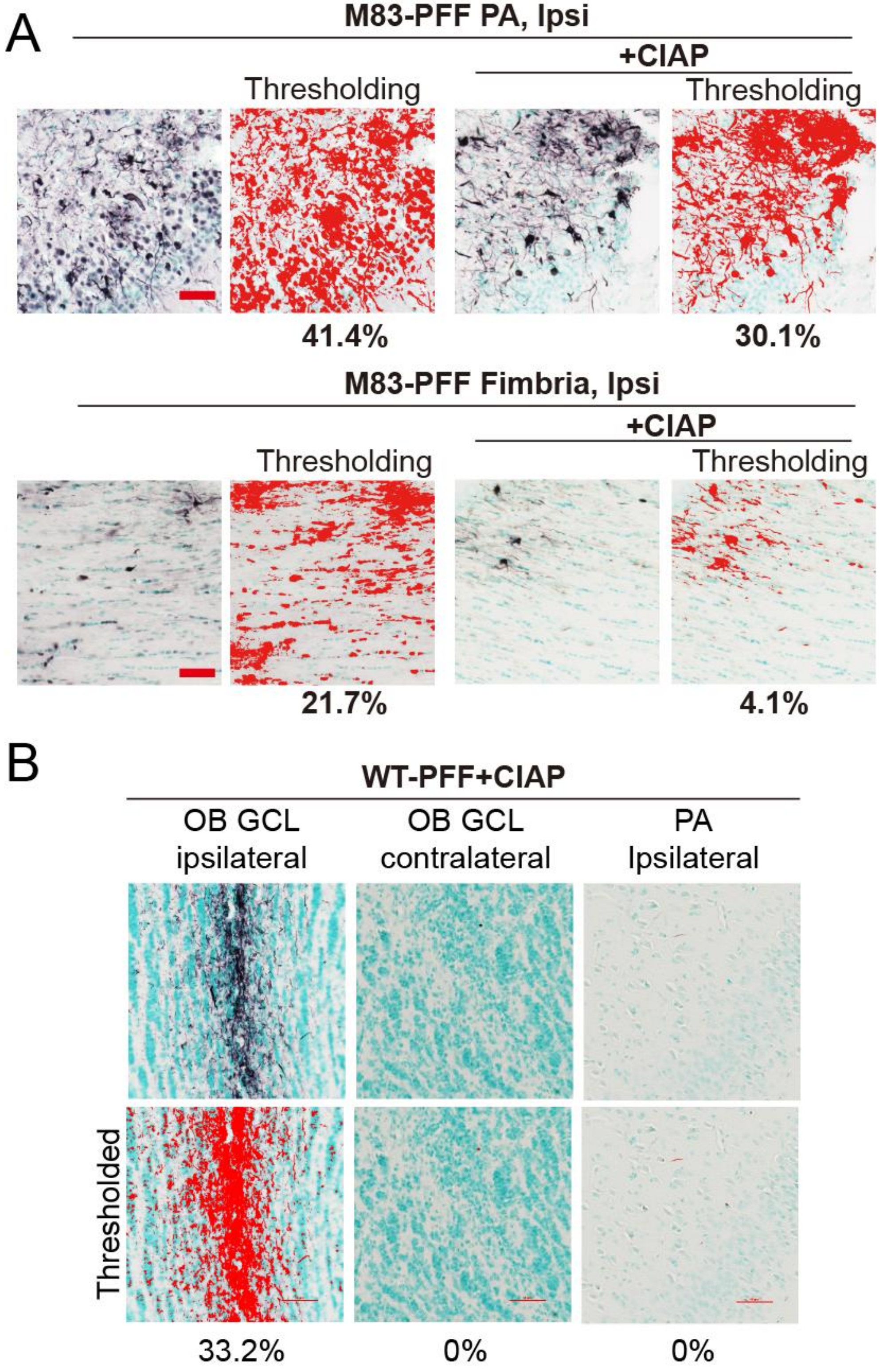


Figure S1. Representative RGB thresholding used to quantify PS129-positive area. PS129-positive area was quantified by RGB thresholding in NIS-Elements software. Threshold parameters were manually optimized to capture dark brown/purple PS129-positive signal while excluding blue methyl green nuclear staining and were then batch-applied to all images. (A) Representative thresholded images from the ipsilateral piriform area (PA) and fimbria of PFF-treated M83 mice, with or without CIAP treatment, are shown with the corresponding percent-area values indicated below each image. (B) Representative images showing PS129 signal quantified in PFF-treated WT mice after CIAP treatment. Scale bars for all images = 50 μm.

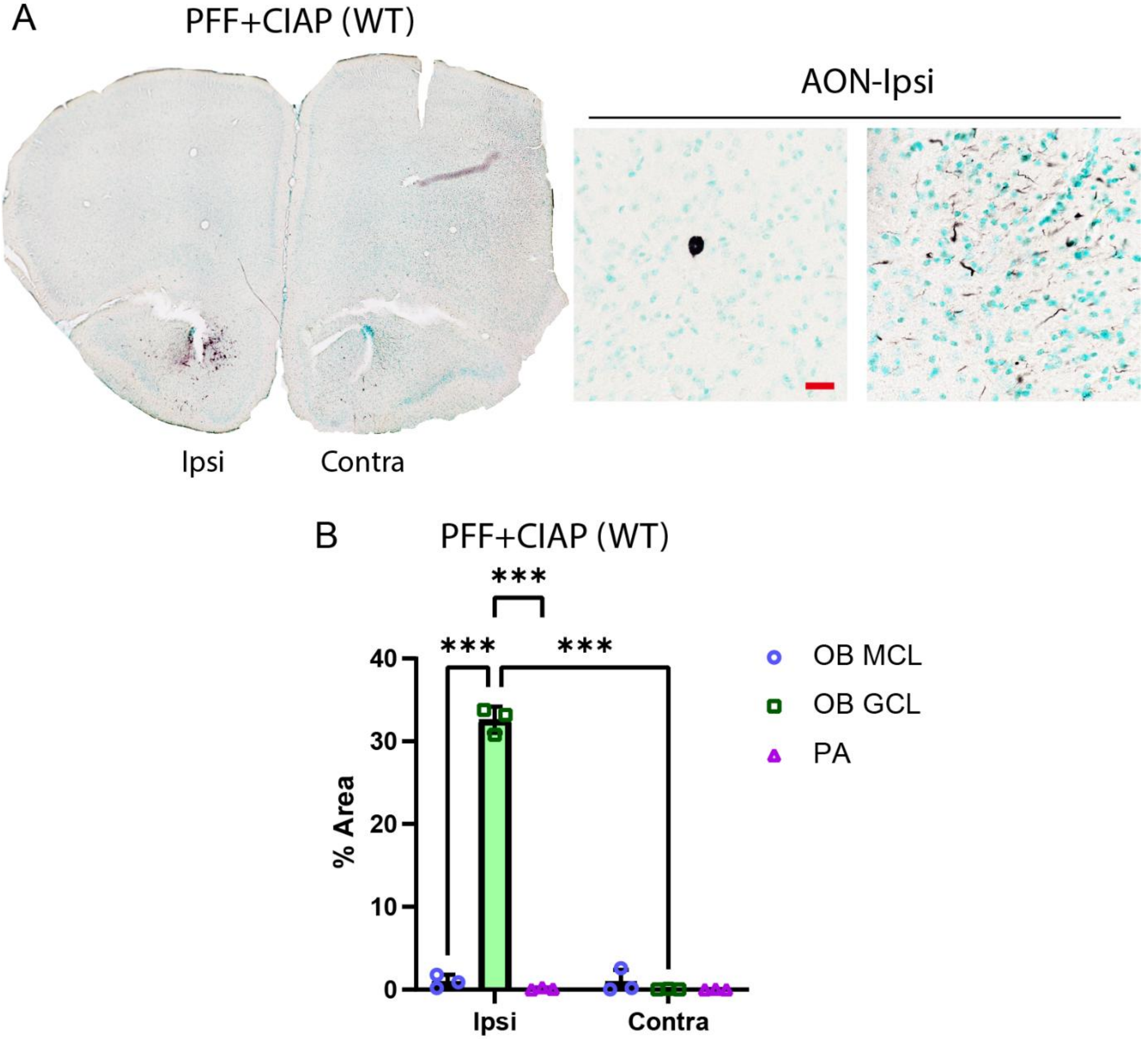


Figure S2. Additional characterization of CIAP-resistant PS129 pathology in the WT-PFF model. (A) Whole-tissue scan of an AON-containing section from a CIAP-treated, PFF-inoculated WT mouse. Representative higher-magnification images of the ipsilateral AON from the same condition are shown alongside. Scale bar = 20 µm. (B) CIAP-resistant PS129-positive area was quantified bilaterally in the OB mitral cell layer (MCL), granule cell layer (GCL), and piriform area (PA) and expressed as percent area. Statistical analysis was performed using two-way ANOVA with Tukey’s multiple comparisons test (***$P < 0.0001$).

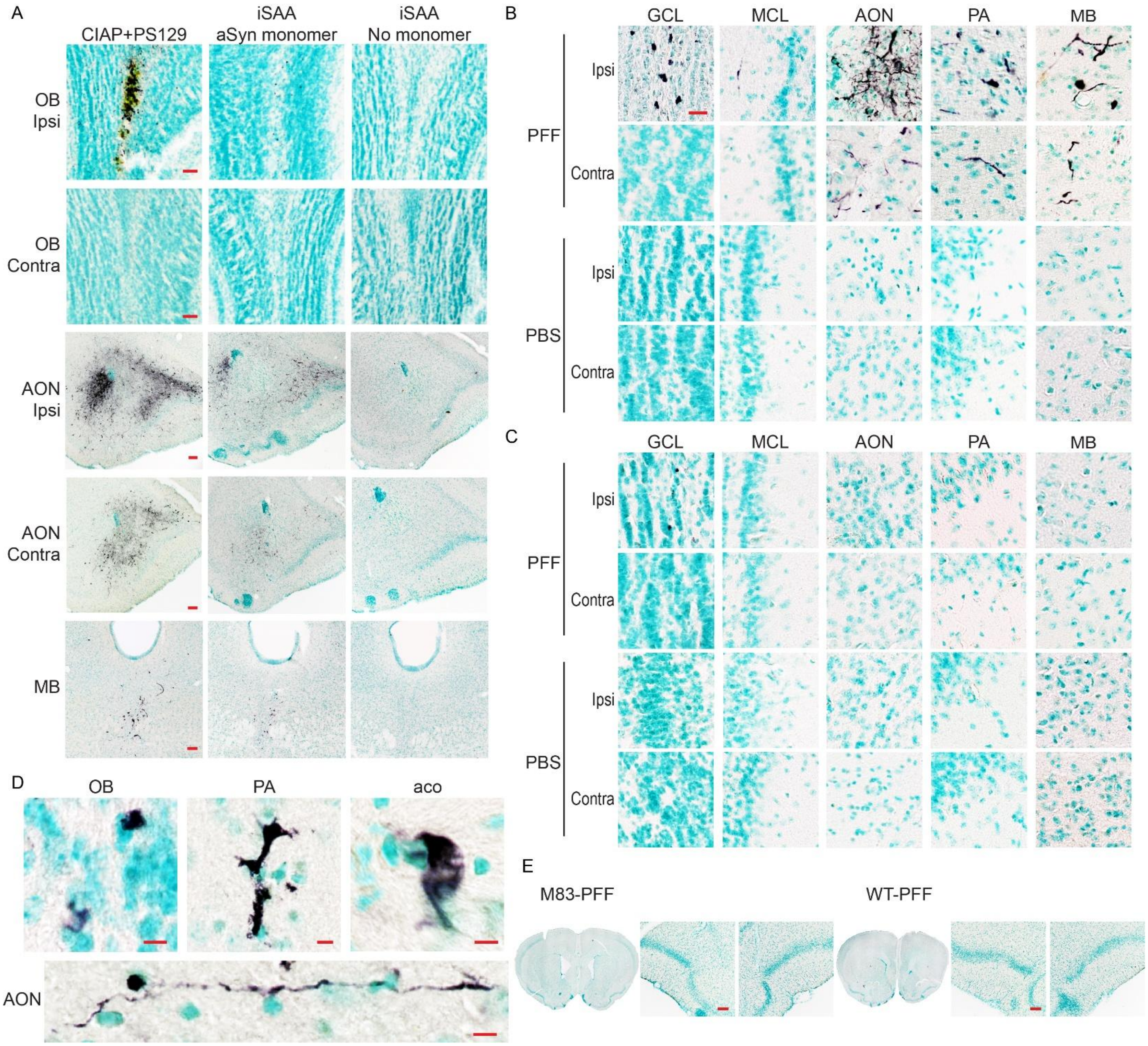


Figure S3. Characterization of iSAA signal in PFF-seeded mouse models. (A) Matched M83-PFF tissue sections were analyzed by CIAP+PS129 staining, iSAA with mouse aSyn monomer, and no-monomer iSAA control. iSAA signal was detected in regions with CIAP-resistant PS129 pathology, including the ipsilateral OB, ipsilateral AON, and MB regions including the periaqueductal gray, but not in the no-monomer control. Scale bars: OB, 50 µm; AON and MB, 100 µm. (B) Higher-magnification iSAA images from both hemispheres of M83-PFF and M83-PBS mice. M83-PFF mice showed stronger ipsilateral signal across examined regions. In the GCL, iSAA-positive structures appeared as variably sized irregular compact deposits with morphology distinct from Lewy bodies. In the AON, prominent dysmorphic neurites with small punctate structures were observed. In the PA and medulla, both intraneuritic and intercellular inclusions were observed. PBS controls showed no detectable signal. Scale bar, 20 µm. (C) iSAA in WT-PFF and WT-PBS mice. WT-PFF mice showed occasional small circular iSAA-positive structures in the GCL, with little to no detectable signal in other regions or PBS controls. Scale bar, 20 µm. (D) High-resolution 60× images showing the morphology of iSAA-positive structures in M83-PFF mice. Signals appeared as irregular compact deposits in the OB, swollen dysmorphic neurites in the PA, crescent-shaped swollen dysmorphic processes in the anterior commissure (aco), and process-associated signal in the AON. Scale bars, 5 μm. (E) Initial iSAA testing using human aSyn monomer in M83-PFF and WT-PFF tissue showed no detectable signal. Scale bars, 20 μm.

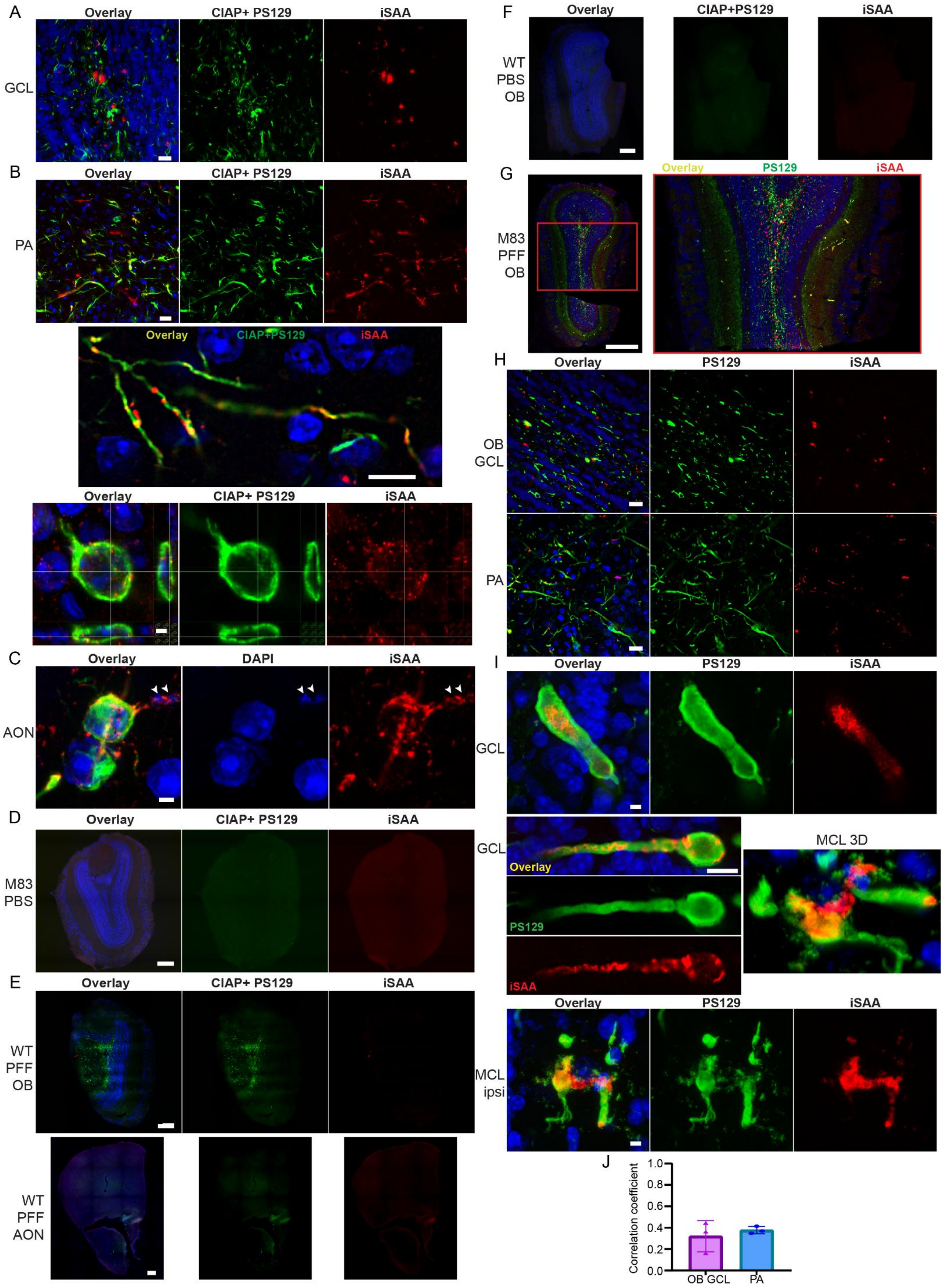

A
Overlay
CIAP+ PS129
iSAA
GCL
B
PA
Overlay
CIAP+PS129
iSAA
C
DAPI
AON
D
M83
PBS
E
WT
PFF
OB
WT
PFF
AON
F
WT
PBS
OB
G
M83
PFF
OB
PS129
H
OB
GCL
I
MCL 3D
MCL
ipsi
J
Correlation coefficient
1.0
0.8
0.6
0.4
0.2
0.0
OB GCL
PA

Figure S4. Multiplex tyramide signal amplification labeling of iSAA-reactive species and CIAP-resistant or total PS129 pathology.

(A-C) Additional images showing dual labeling of iSAA-reactive species and CIAP-resistant PS129 pathology in the ipsilateral hemisphere are shown. (A) 20× magnification image of the OB GCL. Scale bar, 20 μm. (B) PA region shown at higher magnification. First row: The 20× image shows partially overlapping and non-overlapping CIAP-resistant PS129 and iSAA signals in neuritic and Lewy body–like structures. Second row: The 60× images show iSAA signal within or adjacent to CIAP-resistant PS129-positive neurites. Last row: Orthogonal views show a Lewy body–like intracellular inclusion with abundant punctate iSAA signal within and around the CIAP-resistant PS129-positive structure. Scale bars: 20 μm, 10 μm, and 2 μm, respectively. (C) Lewy body–like structure with an associated process showing fragmented DAPI-positive nuclear material within the extended process, with iSAA signal intermingled with or surrounding these fragments. Scale bar, 2 μm. These panels contain additional views and expanded regions, some of which are from the representative image set shown in Figure 3.

(D–F) Dual labeling was performed in PBS-treated M83 mice and WT mice. Ipsilateral hemispheres are shown. Z-stacked images are displayed as maximum-intensity projections. (D) Whole OB scan from an M83 PBS-treated mouse showing no detectable CIAP-resistant PS129 or iSAA signal. Scale bar, 300 μm. (E) Whole-section scans of WT PFF-treated OB and AON showing CIAP-resistant PS129 structures with minimal iSAA signal. Scale bars, 300 μm. (F) Whole OB scan from a WT PBS-treated mouse showing no detectable CIAP-resistant PS129 or iSAA signal. Scale bar, 300 μm.

(G–J) iSAA signal was also dual-labeled with total PS129 in M83 PFF-treated mice, and ipsilateral hemispheres are shown. (G) Whole OB scan shown as a z-stack maximum-intensity projection. The boxed region is enlarged on the right. Scale bar, 500 μm. (H) 20× magnification images of the OB GCL and PA showing total PS129 and iSAA labeling. Scale bars, 20 μm. (I) First row: OB GCL showing swollen PS129-positive neurites containing iSAA deposits. Middle row, left: PS129-positive somatic and neuritic structures containing iSAA signal. Middle row, right: 3D reconstruction of a PS129-positive structure detected around MCL. Last row: single optical plane from the 3D image, showing iSAA signal extending from one neuronal structure toward an adjacent dysmorphic neurite. Scale bars: 5 μm, 10 μm, and 5 μm, respectively. (J) Pearson's correlation analysis showed moderate colocalization between total PS129 and iSAA signals in the OB GCL and PA of M83-PFF mice. Mean Pearson's correlation coefficients were 0.3213 in OB GCL and 0.3771 in PA, with SDs of 0.1461 and 0.03372, respectively.

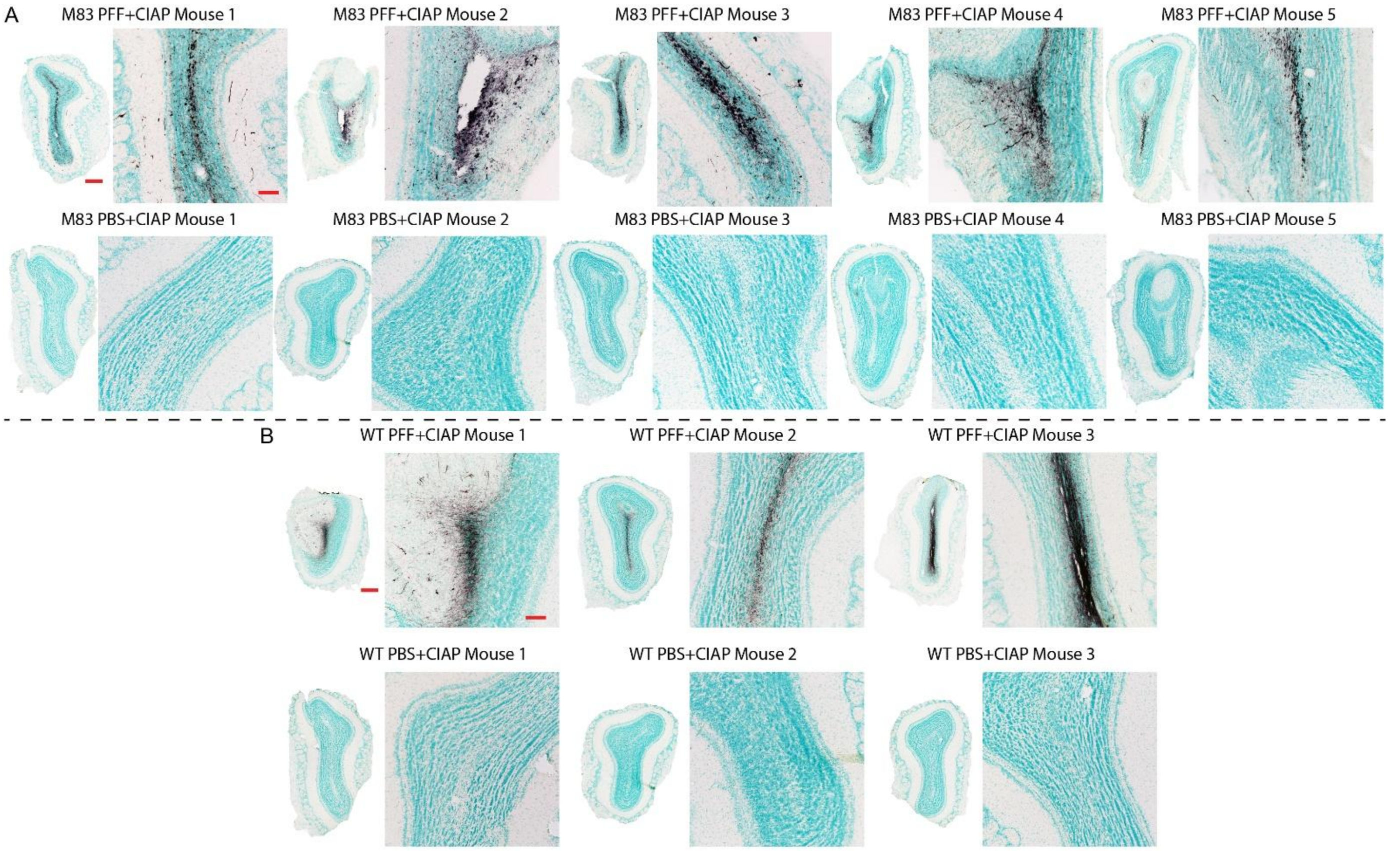


Figure S5. CIAP-resistant PS129 labeling in OB sections used for BAR analysis. Representative OB sections from the cases used for BAR-PS129 interactome mapping are shown after CIAP pretreatment and PS129 immunolabeling. (A) M83 mice treated with PFF or PBS. PFF-treated M83 mice showed robust CIAP-resistant PS129 labeling in the OB, whereas PBS-treated M83 controls showed no residual signal after CIAP treatment. (B) WT mice treated with PFF or PBS. WT PFF-treated mice showed readily detectable CIAP-resistant PS129 labeling at the OB injection site, whereas WT PBS controls showed no detectable CIAP-resistant PS129 signal. For each case, a low-magnification OB overview is shown alongside a higher-magnification image of the corresponding labeled region. Scale bars: whole-tissue scans, 300 μm; higher-magnification images, 100 μm.

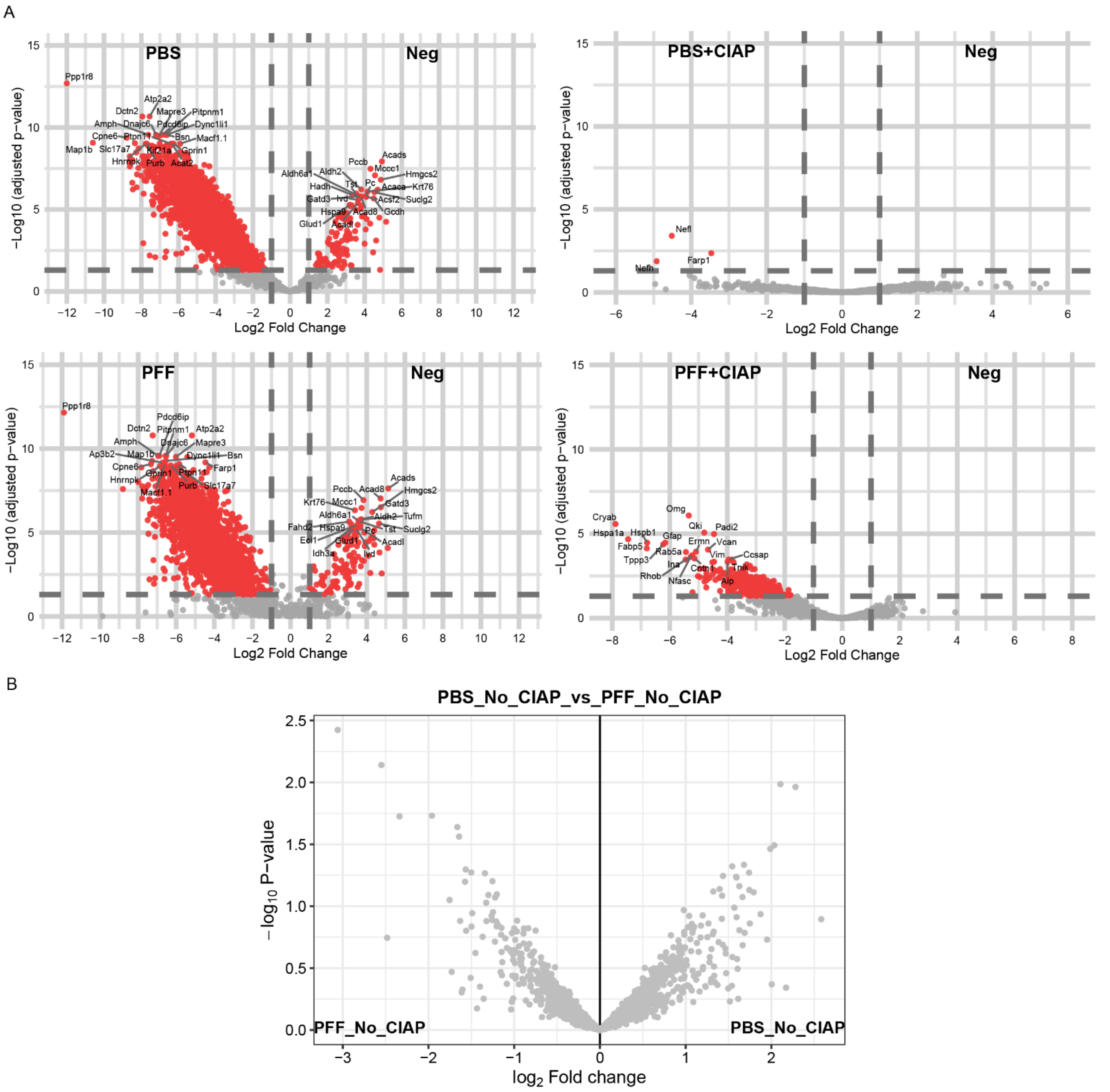


Figure S6. Volcano plot analysis of BAR-PS129–enriched proteins in M83 OB samples. (A) Volcano plots showing differential protein abundance in PBS, PBS+CIAP, PFF, and PFF+CIAP samples relative to their corresponding negative controls (Neg). Significantly enriched proteins are shown in red, and non-significant/background proteins are shown in gray. The top 20 enriched proteins in each comparison are annotated by gene name. (B) LFQ-Analyst-generated volcano plot comparing BAR-PS129–enriched proteins in M83 PBS and M83 PFF samples without CIAP pretreatment. No proteins were significantly different between conditions.

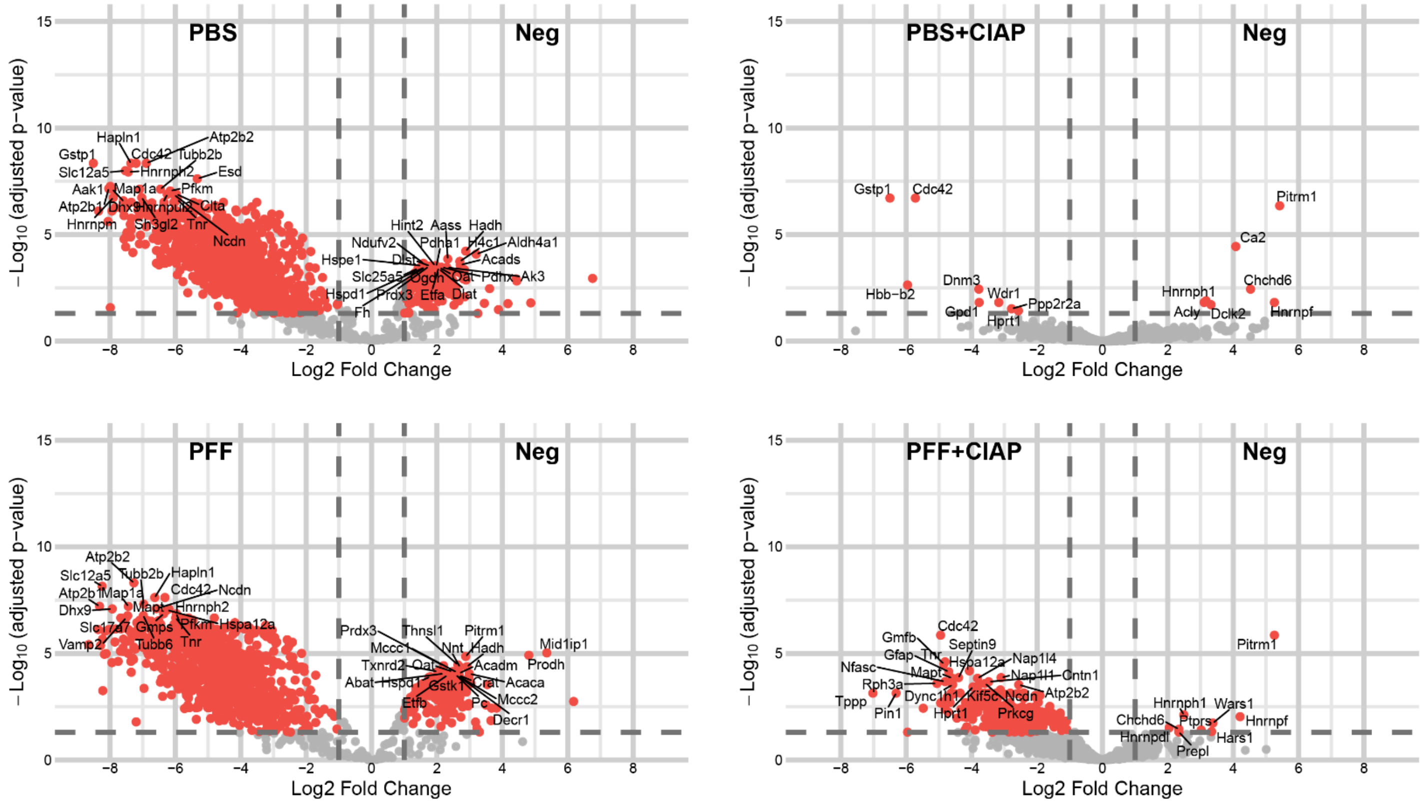


Figure S7. Volcano plot analysis of BAR-PS129 enriched proteins in WT OB samples. (A) Volcano plots showing differential protein abundance in PBS, PBS+CIAP, PFF, and PFF+CIAP samples relative to their corresponding negative controls (Neg). Significantly enriched proteins are shown in red, whereas non-significant/background proteins are shown in gray. The top 20 enriched proteins in each comparison are annotated by gene name.

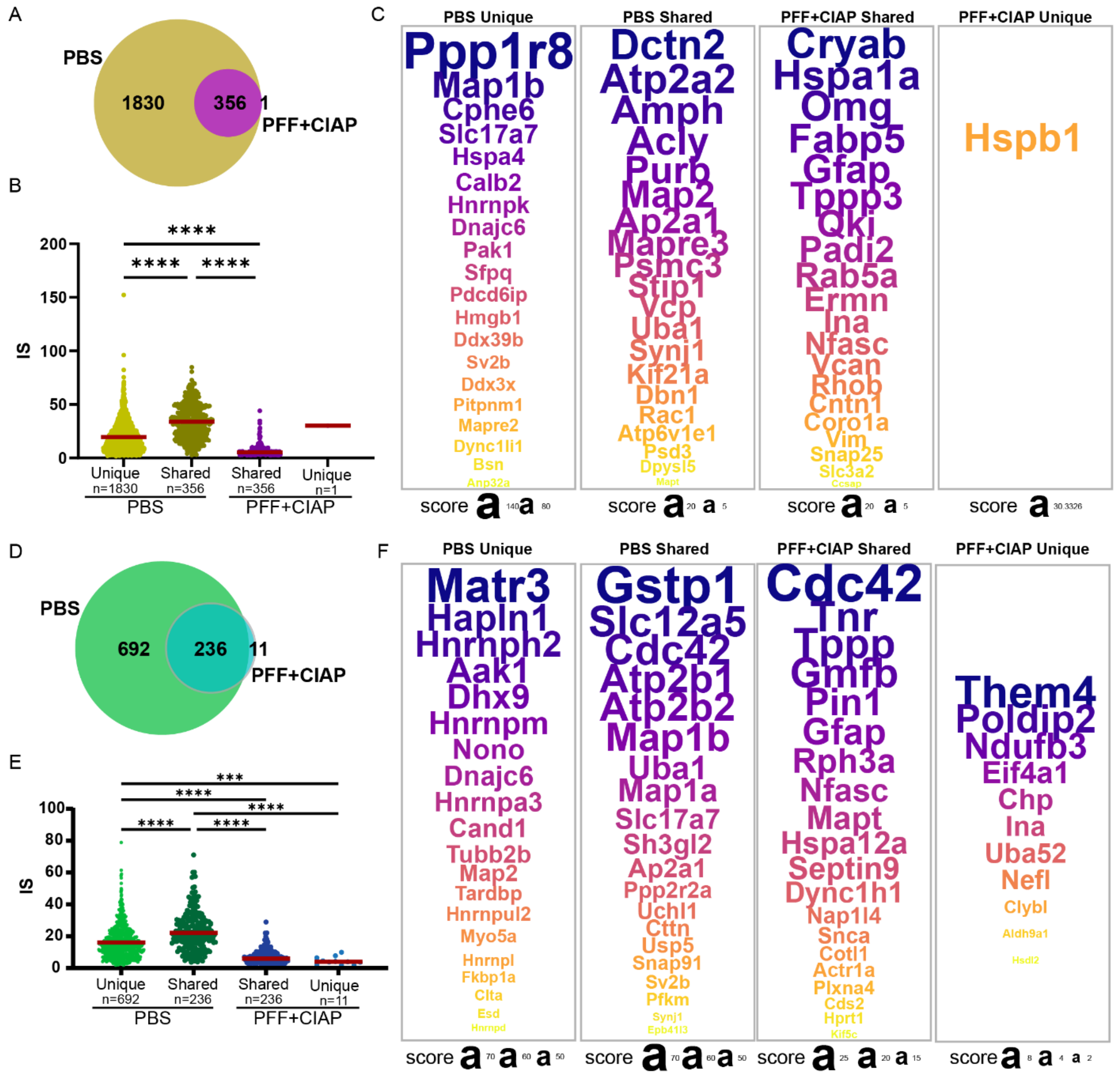


Figure S8. Comparison of PBS and PFF+CIAP PS129 interactomes in M83 and WT mice.
(A) Weighted Venn diagram comparing M83 PBS and PFF+CIAP PS129 interactomes, generated using DeepVenn (arXiv:2210.04597). In M83 mice, 1,830 proteins were unique to PBS, 1 protein was unique to PFF+CIAP, and 356 proteins overlapped between conditions. (B) Proteins were ranked by importance score (IS). PBS-shared proteins had the highest mean IS and were significantly higher than both PBS-unique and PFF+CIAP-shared proteins. PBS-unique proteins also had significantly higher IS than PFF+CIAP-shared proteins. Statistical comparisons were performed using Kruskal–Wallis test followed by Dunn's multiple comparisons test; ****$p < 0.0001$. Mean IS ± SD values were: PBS unique, 19.57 ± 13.76; PBS shared, 33.91 ± 16.24; PFF+CIAP shared, 6.78 ± 4.85. The single PFF+CIAP-unique protein was HSPB1, with an IS of 30.33. (C) Top IS-ranked proteins from each M83 group are shown as word plots, with word size proportional to IS within each group. (D) The same analysis was performed in WT mice. In WT mice, 692 proteins were unique to PBS, 11 proteins were unique to PFF+CIAP, and 236 proteins overlapped between conditions. (E) Proteins were ranked by IS in WT mice. Mean IS ± SD values were: PBS unique, 16.12 ± 10.07; PBS shared, 23.49 ± 12.07; PFF+CIAP shared, 7.09 ± 4.29; and PFF+CIAP unique, 4.57 ± 0.28. (F) Top IS-ranked proteins from each WT group are shown as word plots, with word size proportional to IS within each group.

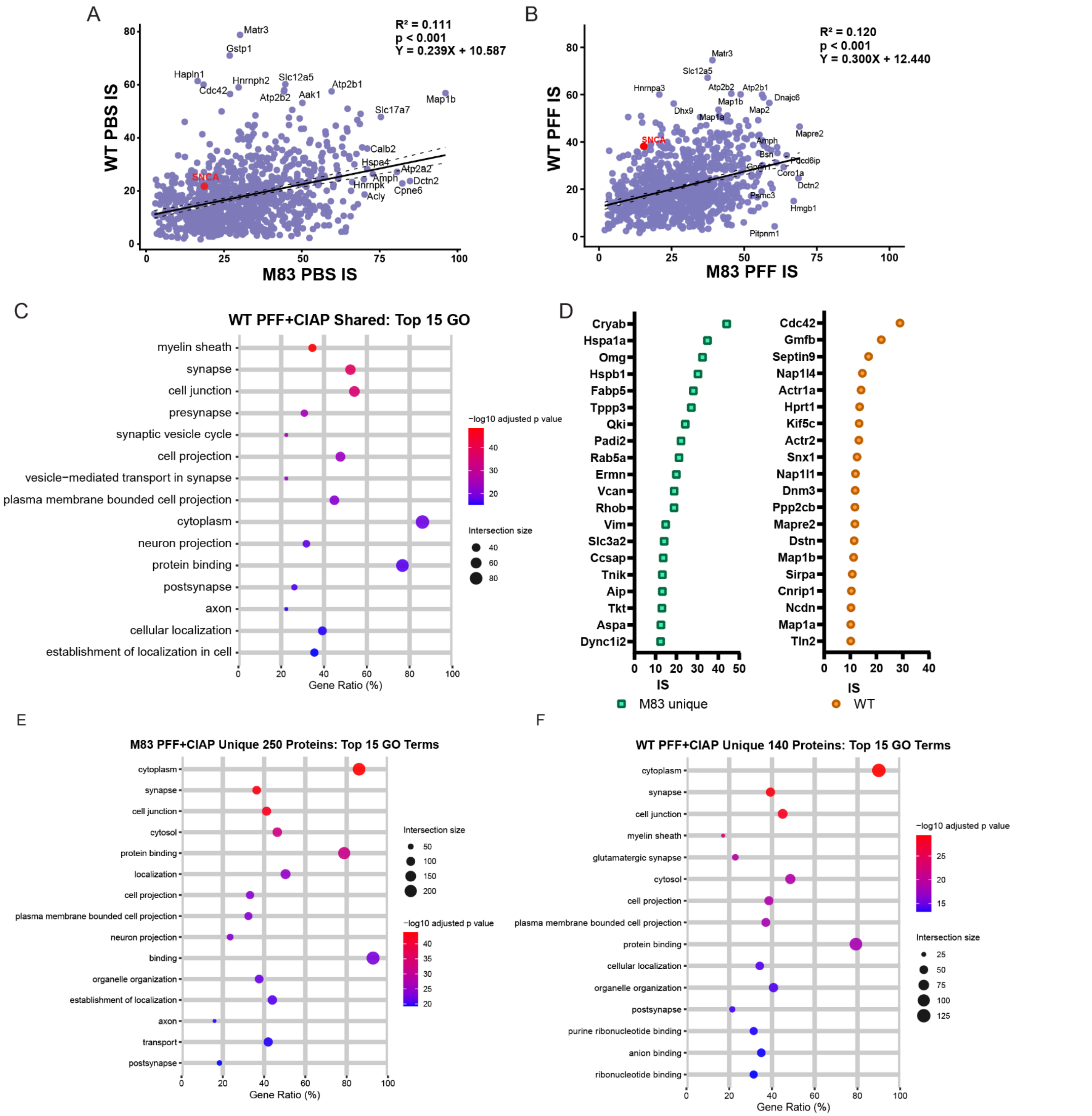


Figure S9. Model-specific and shared features of WT and M83 PS129 interactomes. (A, B) Correlation plots of proteins shared between M83 and WT models under PBS (A) and PFF (B) conditions. Both comparisons showed modest positive correlations between models. SNCA ranked higher in WT than in M83 under both conditions. (C) Top 15 GO terms for the consensus CIAP-resistant PS129 interactome ranked by WT importance score (IS). Myelin sheath was the most significant term, consistent with the corresponding M83-ranked GO analysis. (D) Top 20 IS-ranked proteins from the M83 and WT PFF+CIAP interactomes. (E) GO analysis of the 250 ranked CIAP-resistant PS129-associated proteins unique to M83. The top terms included cytoplasm, synapse, and cell junction, with neuron projection and axon also represented among the top 15 terms. (F) GO analysis of the 140 ranked CIAP-resistant PS129-associated proteins unique to WT. Similar to M83, the top terms included cytoplasm, synapse, and cell junction, whereas myelin sheath, glutamatergic synapse, and ribonucleotide-binding terms were more prominent in WT.

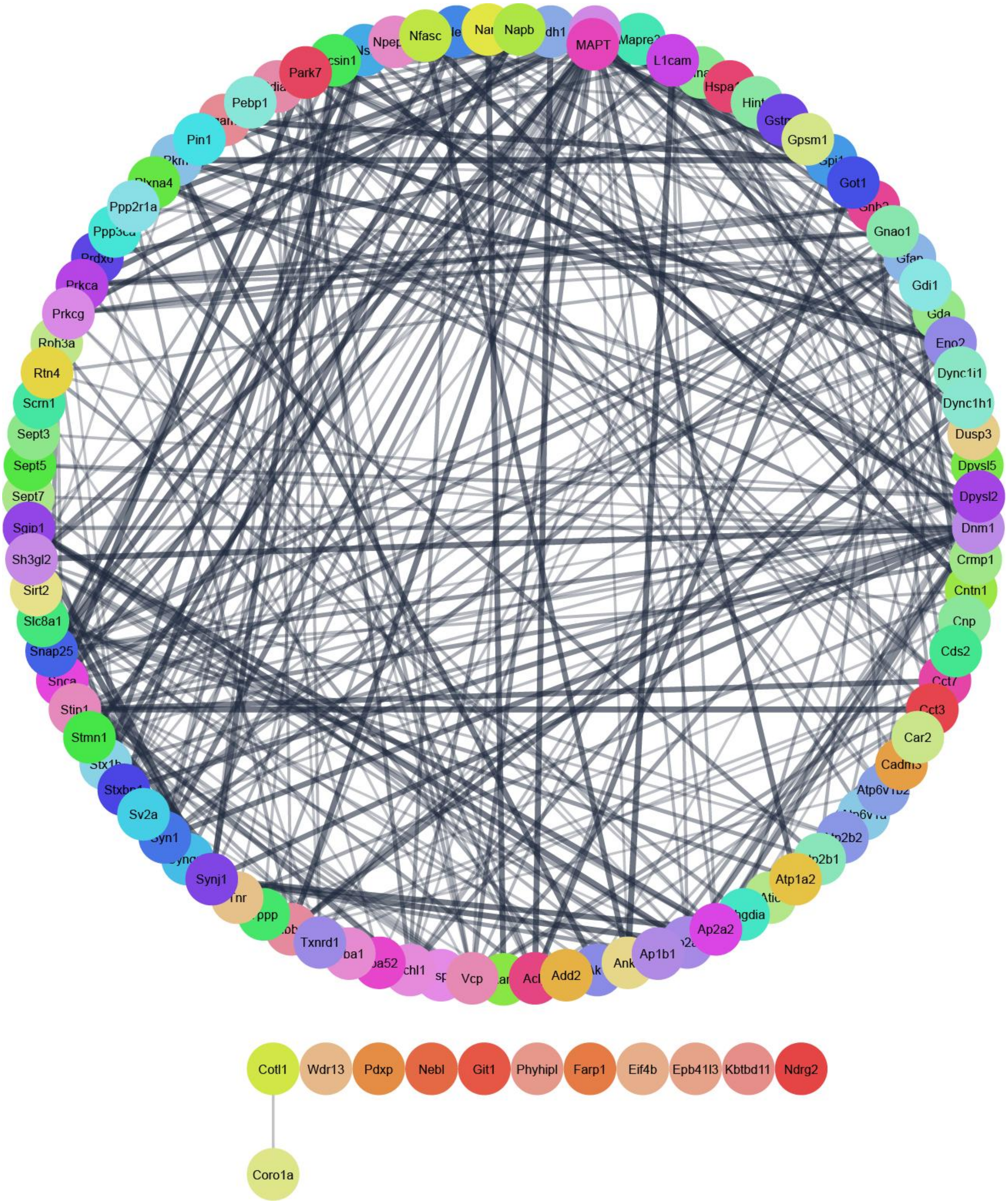


Figure S10. Functional interaction network of the 107 consensus CIAP-resistant PS129-associated proteins. The 107 consensus CIAP-resistant PS129-associated proteins were mapped in Cytoscape STRING to assess functional interactions. A total of 95 proteins formed a main interaction cluster. Of the remaining proteins, 2 formed a doublet and 10 were singletons. Nodes represent proteins, and edges represent functional interactions, with edge thickness reflecting STRING interaction scores. Confidence score cutoff, 0.4.

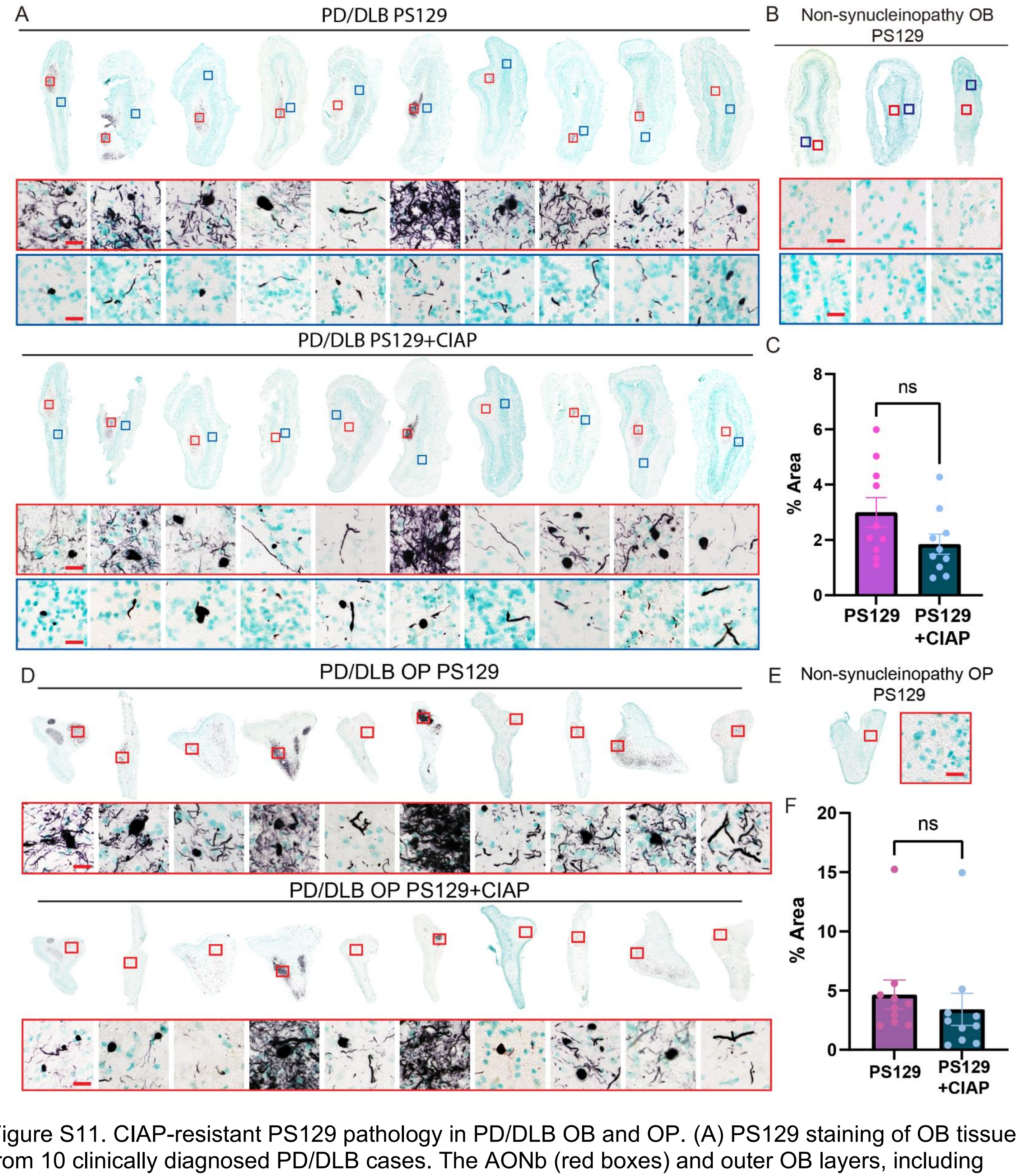


Figure S11. CIAP-resistant PS129 pathology in PD/DLB OB and OP. (A) PS129 staining of OB tissue from 10 clinically diagnosed PD/DLB cases. The AONb (red boxes) and outer OB layers, including the GCL, MCL, and IPL (blue boxes), are enlarged below the whole-tissue scans. PS129-positive structures persisted in matched CIAP-pretreated sections from the same cases. Scale bars = 20 µm. (B) PS129 staining of three non-synucleinopathy OB cases showed no detectable immunoreactivity. Scale bars = 20 µm.

(C) Quantification showed no significant difference in PS129 signal between PD/DLB OB sections with or without CIAP pretreatment. Mean PS129 = 3.0, SD = 1.698; mean PS129 + CIAP = 1.847, SD = 1.145. Two-tailed unpaired t-test; ns, not significant. (D) Olfactory peduncle (OP) sections from the same cases shown in (A) were stained for PS129. OP sections were analyzed without stratification by concentric layer. PS129-positive regions were selected, boxed in red, and enlarged below the corresponding OP images. In matched CIAP-pretreated sections, aggregated PS129 persisted. Scale bars = 20 μm. (E) No PS129 immunoreactivity was observed in OP section from a non-synucleinopathy case. Scale bars = 20 μm. (F) Quantification showed no significant difference in PS129 signal between PD/DLB OP sections with or without CIAP pretreatment. Mean PS129 = 4.666, SD = 3.897; mean PS129 + CIAP = 3.412, SD = 4.294. Two-tailed unpaired t-test; ns, not significant.

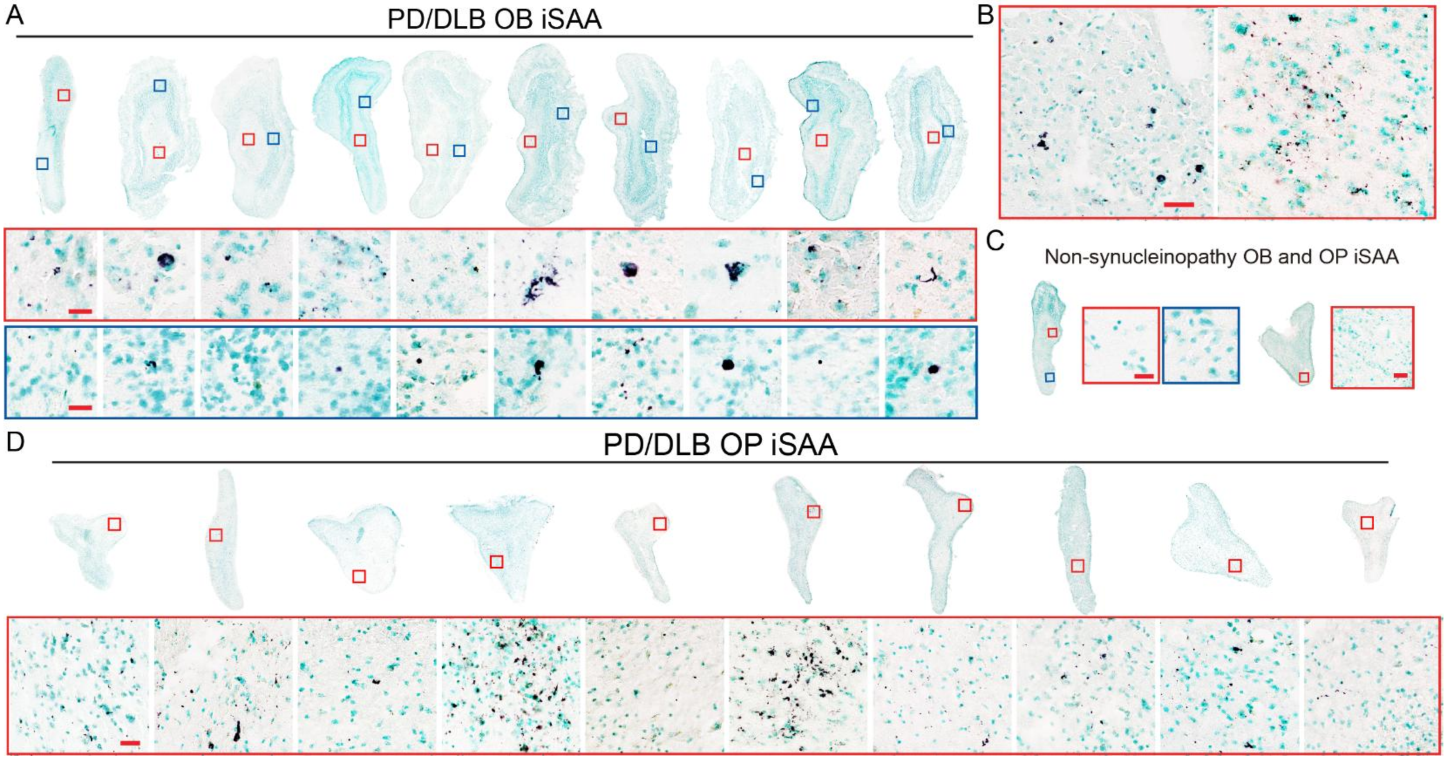


Figure S12. iSAA detection in PD/DLB OB and OP. The same 10 clinically diagnosed PD/DLB cases used for PS129 staining were tested by iSAA. (A) OB sections are shown with higher-magnification images of the AONb (red boxes) and outer OB layers (blue boxes) below each whole-tissue scan. iSAA signal was detected in all cases, although signal intensity varied across cases and was most prominent in the AONb. Scale bars = 20 μm. (B) Representative iSAA signal distribution in the AONb is shown. Scale bars = 50 μm. (C) Non-synucleinopathy OB and OP sections tested by iSAA are shown with corresponding higher-magnification images on the right. OB scale bars = 20 μm; OP scale bars = 50 μm. (D) OP sections from the same 10 PD/DLB cases were tested by iSAA. Because OP sections were analyzed without stratification by concentric layer, regions containing iSAA signal were enlarged and shown below each corresponding OP image. Scale bars = 50 μm.

| # | M83-OB Model Sample ID | (mg) |
|---|---|---|
| 1 | BAR-Neg PFF 1 | 1.1 |
| 2 | BAR-PS129 PFF 1 | 3 |
| 3 | BAR-PS129+CIAP PFF 1 | 3.2 |
| 4 | BAR-Neg PBS 1 | 2.4 |
| 5 | BAR-PS129 PBS 1 | 3.3 |
| 6 | BAR-PS129+CIAP PBS 1 | 4.2 |
| 10 | BAR-Neg PBS 2 | 3.2 |
| 11 | BAR-PS129 PBS 2 | 2.7 |
| 12 | BAR-PS129+CIAP PBS 2 | 7.5 |
| 13 | BAR-Neg PFF 2 | 2.6 |
| 14 | BAR-PS129 PFF 2 | n/a |
| 15 | BAR-PS129+CIAP PFF 2 | 4.1 |
| 16 | BAR-Neg PFF 3 | 1.8 |
| 17 | BAR-PS129 PFF 3 | 1.9 |
| 18 | BAR-PS129+CIAP PFF 3 | 2.5 |
| 19 | BAR-Neg PBS 3 | n/a |
| 20 | BAR-PS129 PBS 3 | 3.5 |
| 21 | BAR-PS129+CIAP PBS 3 | 7.4 |
| 22 | BAR-Neg PBS 4 | n/a |
| 23 | BAR-PS129 PBS 4 | 4.6 |
| 24 | BAR-PS129+CIAP PBS 4 | 6.6 |
| 25 | BAR-Neg PFF 4 | n/a |
| 26 | BAR-PS129 PFF 4 | n/a |
| 27 | BAR-PS129+CIAP PFF 4 | n/a |
| 28 | BAR-Neg PBS 5 | 1.8 |
| 29 | BAR-PS129 PBS 5 | 3 |
| 30 | BAR-PS129+CIAP PBS 5 | 2.7 |
| 31 | BAR-Neg PFF 5 | n/a |
| 32 | BAR-PS129 PFF 5 | 5.9 |
| 33 | BAR-PS129+CIAP PFF 5 | 4.2 |
| | AVERAGE | 3.617391304 |
| | STDEV.S | 1.757232373 |

| # | WT-OB Model Sample ID | (mg) |
|---|---|---|
| 1 | BAR-Neg V1 | 4.1 |
| 2 | BAR-Neg V2 | 3.7 |
| 3 | BAR-Neg V3 | 5.3 |
| 4 | BAR-Neg P1 | 5.3 |
| 5 | BAR-Neg P2 | 5.8 |
| 6 | BAR-Neg P3 | 5.1 |
| 7 | BAR-PS129 V1 | 3.3 |
| 8 | BAR-PS129 V2 | 5.9 |
| 9 | BAR-PS129 V3 | 4.3 |
| 10 | BAR-PS129 P1 | 4.5 |
| 11 | BAR-PS129 P2 | 5 |
| 12 | BAR-PS129 P3 | 4.7 |
| 13 | BAR-PS129+CIAP V1 | 4.2 |
| 14 | BAR-PS129+CIAP V2 | 4 |
| 15 | BAR-PS129+CIAP V3 | 3.4 |
| 16 | BAR-PS129+CIAP P1 | 4.9 |
| 17 | BAR-PS129+CIAP P2 | 4 |
| 18 | BAR-PS129+CIAP P3 | 3.6 |
| | Average | 4.505555556 |
| | STDEV.S | 0.790734746 |

Table S1. Combined wet mass of M83 and WT OB samples used for BAR captures.
